# Transcriptional control of apical protein clustering drives *de novo* cell polarity establishment in the early mouse embryo

**DOI:** 10.1101/2020.02.10.942201

**Authors:** Meng Zhu, Peizhe Wang, Charlotte E. Handford, Jie Na, Magdalena Zernicka-Goetz

## Abstract

The establishment of cell polarity *de novo* in the early mammalian embryo triggers the transition from totipotency to differentiation to generate embryonic and extra-embryonic lineages. However, the molecular mechanisms governing the timing of cell polarity establishment remain unknown. Here, we identify stage-dependent transcription of Tfap2c and Tead4 as well as Rho GTPase signaling as key for the onset of cell polarization. Importantly, advancing their activity can induce precocious cell polarization and ectopic lineage differentiation in a cell-autonomous manner. Moreover, we show that the asymmetric clustering of apical proteins, regulated by Tfap2c-Tead4, and not actomyosin flow, mediates apical protein polarization. These findings identify the long-sought mechanism for the onset of polarization and the first lineage segregation in the mouse embryo.

## Introduction

The fertilized egg is totipotent and so can give rise to any embryonic or extra-embryonic tissue. In the mammalian embryo, totipotency becomes restricted as cells undertake the first cell fate decision and generate two distinct cell populations: the inner cell mass (ICM) and the outer extra-embryonic trophectoderm (TE). The ICM will form the epiblast (EPI), the future fetus, and the extra-embryonic primitive endoderm (PE), the future yolk sac. The TE will form the placenta. Formation of these three cell lineages by the time of implantation is a prerequisite for a successful pregnancy. Embryo polarization is key to the segregation of the ICM and TE lineages (Johnson and Ziomek, 1981a; Korotkevich et al., 2017) and in the mouse, this process happens at the late 8-cell stage(Fleming et al., 1986; Johnson and Ziomek, 1981a, b) when all cells acquire apical domains composed of the conserved Par protein complex, ERM proteins and enclosed by an actomyosin ring (Louvet et al., 1996; Plusa et al., 2005). In subsequent cell divisions, cells that retain this apical domain express the transcription factors Cdx2 and Gata3 to acquire TE fate, whereas the apolar cells maintain pluripotency to become ICM (Nishioka et al., 2009; Ralston et al., 2010).

Despite the importance of cell polarization for lineage segregation, the mechanisms establishing cell polarization and its timing in the mammalian embryo remain unknown. The apical domain present at the late 8-cell stage is unique as the ability of the cells to polarize is temporally restricted specifically to this stage. In addition, the apical domain can be established in a self-organized manner and in the absence of cell adhesion. However, the identity of the upstream genetic regulators and the mechanism driving cell polarity establishment remain unknown. Here, we use a combination of embryological, gene-editing and live-imaging methods to identify the crucial factors sufficient to trigger apical domain assembly, and the mechanisms by which these factors act.

## Results

### Transcription is required for apical domain assembly

It is well established that mouse embryo blastomeres polarize at the late 8-cell stage. We have recently found that apical domain formation requires actomyosin activation downstream of Rho GTPase signalling (Zhu et al., 2017). However, Rho signalling alone is insufficient to advance the timing of cell polarization (Zhu et al., 2017), suggesting the existence of additional factors parallel to actomyosin for apical domain formation. As there appeared to be a relationship between the developmental timing of zygotic genome activation and the onset of cell polarity establishment in different mammalian species(Brunet-Simon et al., 2001; Graf et al., 2014; Koyama et al., 1994; Nikas et al., 1996; Telford et al., 1990), we hypothesized that zygotic transcription might play a key regulatory role in this process. To address this hypothesis, we deployed two complementary approaches: first, to inhibit transcription prior to cell polarization and second, to increase the concentration of zygotic transcripts. In both cases, we examined the impact of these manipulations on the timing of apical domain formation.

To inhibit transcription, we treated embryos from the early 8-cell stage with two different transcription inhibitors, 5,6-Dichlorobenzimidazole 1-β-D-ribofuranoside DRB (Bensaude, 2011; Efrat and Kaempfer, 1984) and Triptolide (Bensaude, 2011; Vispe et al., 2009) and examined apical protein localization (Fig. 1A-D; Fig. S1A-G). Inhibition of transcription with either drug led to the failure of apical positioning of the polarity marker, Pard6 (Fig. 1B-D; Fig. S1C-E) although cells continued to undergo cytokinesis (Fig. S1A-B; F-G). Washing-out the reversible transcription inhibitor DRB allowed the resumption of apical domain formation after 9 h, suggesting that failure to polarize was a consequence of transcriptional inhibition (Fig. S1H-J). Thus, transcription is required from the early 8-cell stage for the mouse embryo to polarize.

**Figure 1.**
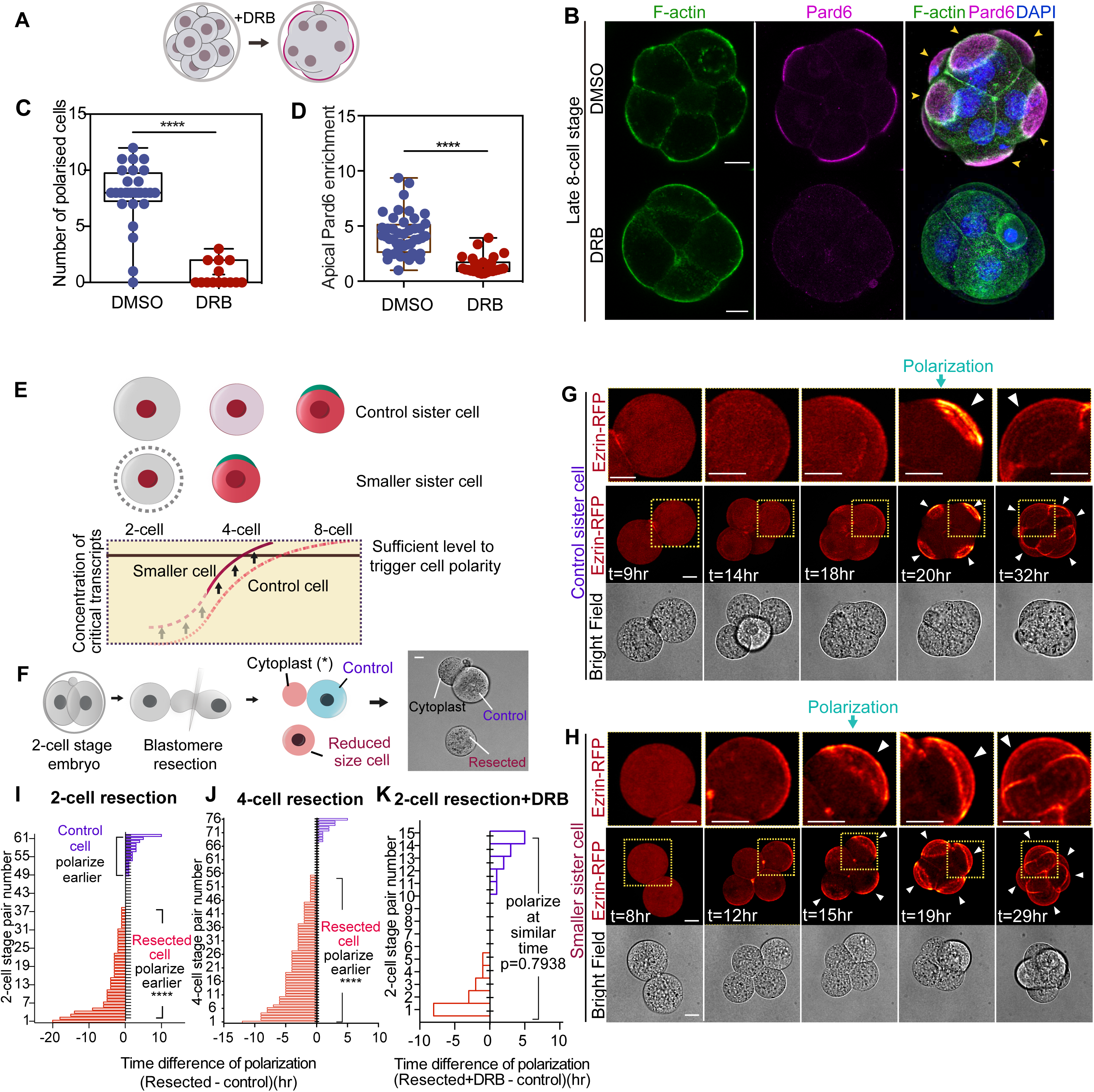
The dependency of cell polarity on nascent transcripts. **(A)** Scheme indicating the regime of treatment with transcription inhibitors DRB or Triptolide from early 8-cell to 8-16 cell stages. (**B**) DMSO (control)- or DRB-treated embryos analyzed at 8-16 cell stage for F-actin, Pard6 and DNA. (**C**) Quantifications of number of polarized cells at the late 8-cell stage in embryos treated with DMSO (control) or DRB. ****p<0.0001, Mann-Whitney test. Each dot represents an analyzed embryo. N= 26 embryos for DMSO, N=14 embryos for DRB. N=2 independent experiments. (**D**) The extent of cell polarization in each cell quantified by the intensity of apical enrichment of Pard6 (see Methods) in cells treated with DMSO (control) or DRB at the 8-16 cell stage. ****p<0.0001. Mann-Whitney test. Each dot represents an analyzed cell. (**E**) Schematic representation of the hypothesis postulating that newly synthesized factors important for cell polarization accumulate up to a point upon which polarization is induced at the 8-cell stage. Decreasing the cell size would elevate the concentration of such factors leading to an advance in the timing of polarization. **(F)** Schematic diagram of blastomere resection at the 2-cell stage resulting in one cell of reduced size and an attached control sister cell (and cytoplast). (**G,H**) Time-lapse movies of control or sister cell with reduced size from the experiment illustrated in **F**. (**I**) Bar chart showing comparisons of the time difference of polarization between small and control sister cells from the experiment illustrated in **F**, each bar represents one comparison. Cells with reduced size polarize earlier in the significant majority of cases (61.3% of the small cells polarize earlier than their control sister cells, 16.1% of small cells polarize with and 22.6%, later than their sisters, N=62 pairs analyzed). The time difference in hours (hr) is calculated by subtracting the time of polarization of the control cell from the time of polarization of the small cell. ****p<0.0001, one-sample t-test, hypothetical mean =0. N=13 independent experiments. (**J**) Bar chart showing time difference of polarization between small and control sister cells from experiment in Fig. S2F, each bar represents one comparison. Cells with reduced size polarize earlier in 81.2% of cases than their control sister cells. N=76 pairs analyzed. ****p<0.0001, one-sample t-test, hypothetical mean=0. N=6 independent experiments. Representative images are shown in Fig. S2G-H. (**K**) Bar chart showing time difference between control sister cells and smaller DRB-treated cells, from experiments illustrated in Fig. S2I. Each bar represents one comparison. Representative images are shown in Fig. S2J-K. time difference (hr) is calculated by subtracting the time of polarization of the control cell from the time of polarization of the small cell, the same holds true for all resection related experiments. Pulsed inhibition of transcription prevents the early polarization of the cells of reduced size. N=15 pairs analyzed. N=3 independent experiments. ns, not significant, one-sample t-test, hypothetical mean =0. Arrows indicate the apical domains. Scale bars, 15µm.

To increase the concentration of zygotic transcripts, we wished to reduce the cytoplasmic volume as this has been shown to result in an increased concentration of newly synthesized mRNA (Bao et al., 2017; Padovan-Merhar et al., 2015). To this end, we resected 30-40% of cytoplasm from one of the two blastomeres, using a method that has been demonstrated not to compromise developmental potential (Zernicka-Goetz, 1998) (Fig. 1F; Fig. S2A). This resection resulted in a higher concentration of newly transcribed mRNAs in the cytoplasm as assessed by single-molecule fluorescence in situ hybridization (smFISH) (Wang et al., 2012) of a zygotically expressed house-keeping gene, Polr2a (Fig. S2B-E). To determine if blastomere resection affected the timing of cell polarization, we injected embryos with Ezrin-RFP mRNA, to visualize the apical domain in living cells, and then resected blastomeres at either the 2-cell stage or 4-cell stage (Fig. 1F-I; Fig.S2F-H). Both experimental and control embryos established all three lineages at the blastocyst stage, indicating that this procedure did not impair normal development (Fig. S2I-K). Importantly, blastomere resection advanced the timing of cell polarization by 2.1 hr (when carried out at the 2-cell stage; Fig. 1F-I; N=62 pairs; Movie S1) and by 3.3 hr (when carried out at the 4-cell stage; N=76 pairs; Fig. S3F-H; Movie S2). To determine whether the effect on cell polarization we observed was indeed due to transcription, we inhibited transcription in the resected cell, by subjecting it to a 3 hr pulse of DRB. In this case, both resected and control cells polarized simultaneously (Fig. 1K; Fig. S2I-K). Thus, although we cannot exclude the possibility that resection has several effects upon the cell, these results indicate that *de novo* transcription contributes significantly to apical domain formation.

### Redundancy of Tfap2c and Tead4 activity in apical domain formation

We next considered whether zygotic genome activation might either directly activate expression of essential cytoskeletal regulators of cell polarization or indirectly through a specific transcriptional hierarchy. To identify candidate proteins for such regulatory roles, we interrogated previously published single-cell RNA-sequencing data (Goolam et al., 2016) and selected 118 polarity regulators whose transcript levels are upregulated between the 2-cell and 8-cell stages. We also identified 15 transcription factors most likely to be active at this time by analyzing ATAC-seq data (Wu et al., 2016) and of these, we selected 6 that become up-regulated by the 8-cell stage (Fig. S3; Table S1,2). We then downregulated the expression of each of these 124 genes by RNAi and scored the effect on the timing of apical domain formation by time-lapse imaging. We found that depletion of only two of these gene products, the transcription factors Tfap2c or Tead4, prevented cell polarization at the 8-cell stage (Fig. 2A-C, E; Fig. S4A-E). When Tfap2c and Tead4 were depleted individually, polarization was delayed to the 16-cell stage (Fig. 2A-C, E-F) but when both factors were depleted together, cell polarization was abolished (Fig. 2D-H). The depletion of Tfap2c and Tead4 also overcame the precocious cell polarization effect resulting from the reduction of cell volume by blastomere resection. This suggests that Tfap2c and Tead4 mediate premature cell polarization in the resection experiment (Fig. S4F-G).

**Figure 2.**
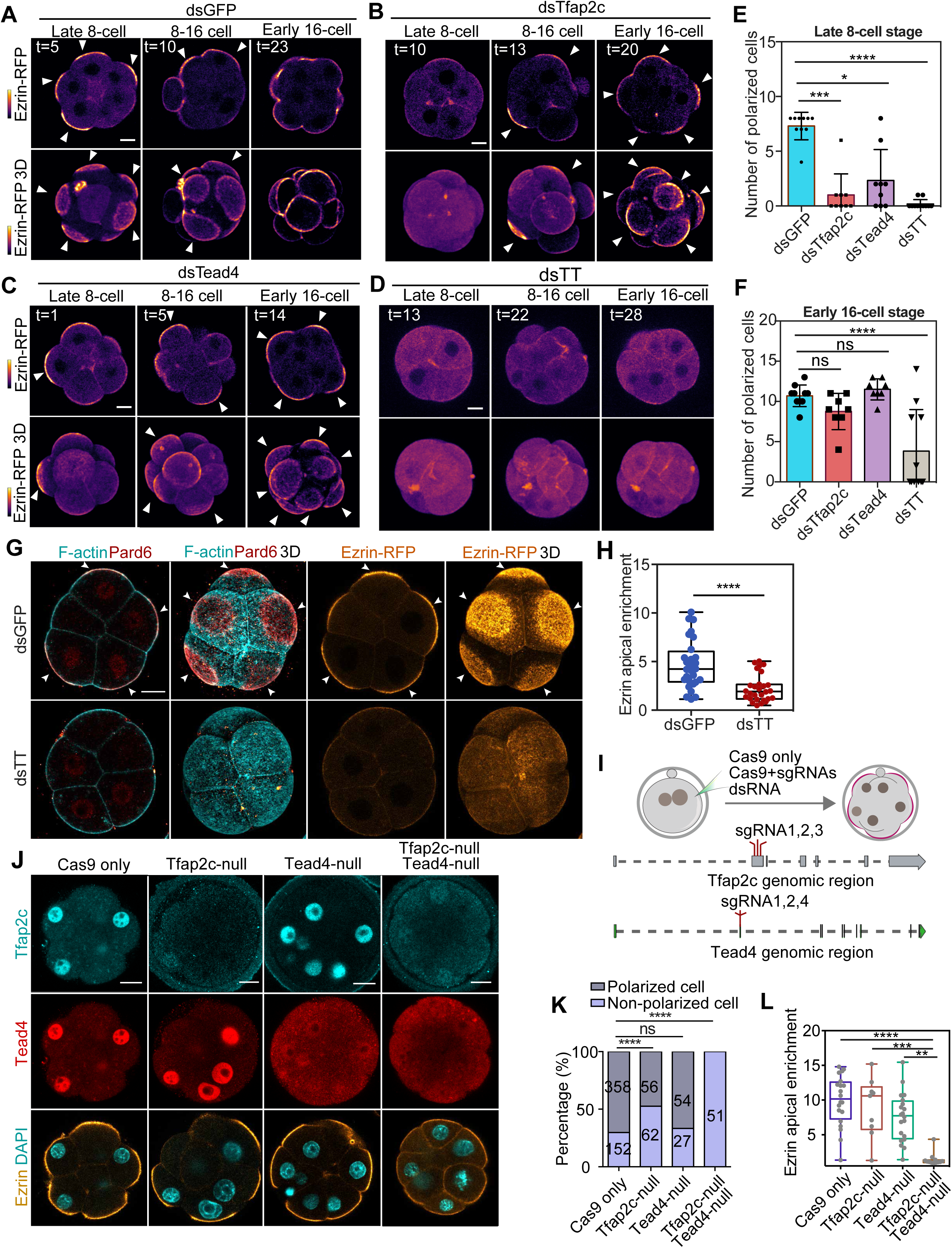
Zygotic expression of Tfap2c and Tead4 is essential for cell polarization. (**A-D**) Time-lapse imaging of embryos injected with Ezrin-RFP mRNA and dsRNAs against GFP; Tfap2c; Tead4; or a mixture of Tfap2c+Tead4 to reveal cell polarization (assessed by Ezrin-RFP localization). Depletion of both Tfap2c and Tead4 leads to failure of polarization until the 16-cell stage. Arrows indicate apical domains. (**E**) Quantification of polarized cell number in late 8-cell stage embryos injected with indicated dsRNAs. N=80 cells (10 embryos) for dsGFP; N=72 cells (9 embryos) for dsTfap2c; N=72 cells (9 embryos) for dsTead4; N=88 cells (11 embryos) for dsTfap2c+dsTead4 (dsTT). *p=0.0306; ***p=0.0006; ****p<0.0001. Kruskal-Wallis test. Data presented as means ± S.D. N=2 independent experiments. (**F**) Quantification of the number of polarized cells in embryos injected with indicated dsRNAs and assessed at the early 16-cell stage. ns, not significant; ****p<0.0001, ordinary one-way ANOVA test. Data presented as means ± S.D. N=2 independent experiments. (**G**) Embryos injected with the dsRNA targeting GFP (control) or Tfap2c and Tead4 and examined at the late 8-cell stage for F-actin, Pard6, and Ezrin localization. White arrows indicate apical domains. (**H**) The extent of polarization in each cell is quantified by the intensity of apical enrichment of Ezrin, see Methods. ****p<0.0001, Student’s t-test. Data are shown as individual data points with Box and Whisker plots (lower: 25%; upper: 75%; line: median; whiskers: min to max). Each dot represents an analyzed cell. (**I**) Schematic of CRISPR-Cas9 strategy used to deplete Tfap2c and Tead4. (**J**) Embryos expressing Ezrin-RFP with wild-type (injected with only Cas9 mRNA, control); single depletion of Tfap2c (injected with Cas9 and sgRNAs targeting only Tfap2c); single depletion of Tead4 (injected with Cas9 and sgRNAs targeting only Tead4); double depletion of Tfap2c and Tead4 (injected with Cas9 and sgRNAs targeting both Tfap2c and Tead4) imaged at 8-16 cell stage to reveal Tfap2c and Tead4 protein levels. (**K**) Proportions of polarized cells in different genotypes presented in **J**. Data presented as a contingency table. Number of cells analyzed presented within each bar. ns, not significant; ****p<0.0001, Fisher’s exact test. N=2 independent experiments. Cells showing Ezrin apical enrichment are polarized (quantification of Ezrin apical/basal signal intensity ratio displayed in **L**). Co-depletion of Tfap2c and Tead4 represses apical domain formation. (**L**) Quantifications of Ezrin apical enrichment in Cas9 only (control), Tfap2c-null, Tead4-null and Tfap2c- and Tead4-null cells. Data are shown as individual data points with Box and Whisker plots (lower: 25%; upper: 75%; line: median; whiskers: min to max). N=20 cells for Cas9 only, N=9 cells for Tfap2c-null, N=20 cells for Tead4-null and N=12 cells for Tfap2c and Tead4-null. Ezrin apical enrichment is calculated as the Ezrin signal intensity on the cell-contact free surface against the Ezrin signal intensity on cell-contacts. **p=0.0012, ***p=0.0007, ****p<0.0001, Kruskal-Wallis test. N=4 independent experiments. Embryos depleted with Tfap2c and/or Tead4 by dsRNA or CRISPR did not show obvious developmental delay. Scale bars, 15µm.

To confirm the roles of Tfap2c and Tead4 in regulating cell polarization, we genetically depleted both genes by CRISPR-Cas9 mutagenesis. We designed three sgRNAs to target a single protein-coding exon of each gene (Fig. 2I) and injected them into the zygote together with Cas9 mRNA and Ezrin-RFP mRNA, to visualize apical domain formation *in vivo*. We then categorized the resultant blastomeres based on whether they had undetectable, moderate, or wild-type levels of Tfap2c or Tead4 proteins at the 8-16 cell stage (Fig. S5A-B). Through DNA sequencing we confirmed that the blastomeres with undetectable Tfap2c or Tead4 were homozygous mutants (subsequently termed Tfap2c-null or Tead4-null) (Fig. 2J; Fig. S5C-D). Simultaneous deletion of *Tfap2c* and *Tead4* completely abolished cell polarization, whereas the effects of their individual deletions were less severe (Fig. 2J-l; Fig. S5E-F), in agreement with our RNAi results above (Fig. 2G-H). These findings lead us to conclude that zygotic expression of *Tfap2c* and *Tead4* is required for cell polarization at the 8-cell stage.

Our findings that Tead4 was involved in the onset of cell polarization were unexpected as thus far, Tead4 has been only known to function downstream of cell polarization, following the nuclear re-localization of its transcriptional co-activator Yap to induce expression of TE transcription factors (Nishioka et al., 2009). To gain further insight into the earlier role of Tead4 that we had now uncovered, we examined the localization of Yap. Surprisingly, we found that Yap localizes to the nucleus together with Tead4 before cell polarization at the 8-cell stage (Fig. S6A-C)(Hirate et al., 2015). This nuclear localization of Yap was diminished by downregulation (Fig. S6D-E), and enhanced by upregulation, of Tead4 expression (Fig. S6F-G). These results suggest that at these earlier stages, Tead4 affects the localization of Yap indicating a polarity-independent Tead4 function.

### Advancing expression of Tfap2c, Tead4 and Rho GTPase induces premature cell polarization

We next wished to determine whether advancing expression of Tfap2c and Tead4 was sufficient to advance the timing of cell polarization. To this end, we injected Tfap2c and Tead4 mRNAs into one blastomere at the 2-cell stage to elevate their expression by the 4-cell stage, together with Ezrin-RFP as an apical marker (Fig. S7A-C). Advancing the expression of Tead4 alone had no obvious effect on cell polarization (Fig. S7D-E,I-J). By contrast, advancing the expression of Tfap2c led to formation of cell protrusions that were enriched in apical polarity proteins, including Pard6 and Ezrin at the late 4-cell stage (Fig. S7F-I). Advancing the expression of Tfap2c and Tead4 together also induced premature formation of cell protrusions (Fig. S7D-G; Fig. 3B-C; Movie S3-4). These Tfap2c- or Tfap2c-Tead4-induced membrane protrusions were smaller than the natural apical domains formed at the late 8-cell stage and lacked the actomyosin-enclosed apical domain that normally forms (Fig. S7H). These results suggested that Tfap2c and Tead4 expression might be sufficient to lead to the polarization of apical proteins but not their expansion to form an actomyosin enclosed cap-like domain.

**Figure 3.**
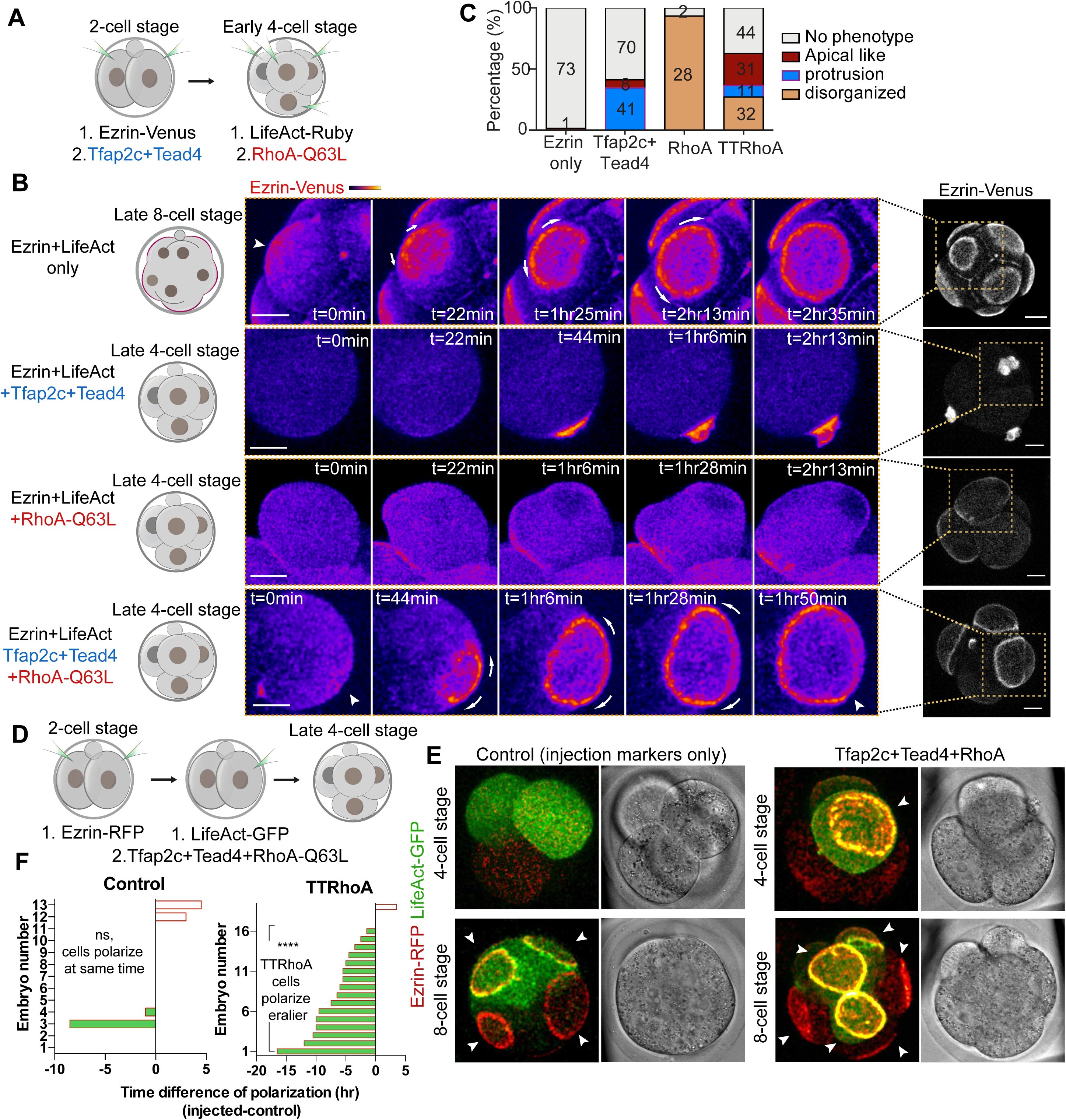
Premature expression of Tfap2c, Tead4, and activated RhoA is sufficient to advance the timing of polarization and differentiation. (**A**) Scheme of Tfap2c, Tead4, and RhoA-Q63L overexpression. (**B**) Time-lapse movies of Ezrin-Venus dynamics during the formation of an apical domain in 1) embryos at late 8-cell stage (from Ezrin-Venus only control group); 2) cells overexpressing Tfap2c+Tead4 showing induced apical protrusions at the late 4-cell stage; 3) cells overexpressing RhoA-Q63L showing disorganized cell morphology at the late 4-cell stage; 4) cells overexpressing Tfap2c+Tead4+RhoA-Q63L showing induced premature full apical domain at the late 4-cell stage. In all conditions the cell divisions were not obviously affected. Short arrows indicate Ezrin-Venus enrichment at the cell-contact free surface. Long arrows indicate the direction of expansion of the Ezrin-Venus enclosed apical domain. Squares indicate the magnified region of each embryo. (**C**) Quantification of structures induced by conditions presented in **B**. Data presented as a stacked bar graph where numbers in each bar indicate the number of cells analyzed. (**D**) Scheme of Tfap2c, Tead4, and RhoA-Q63L overexpression in half of the embryos. (**E**) Representative images of embryos overexpressed with Ezrin-RFP and LifeAct-GFP mRNA only (control) or with Tfap2c, Tead4 and RhoA-Q63L mRNA at 4 -or 8-cell stage. The cells with overexpressed Tfap2c, Tead4 and RhoA-Q63L polarize significantly earlier than control cells in the same embryos, or cells in the control embryos. Arrows indicate the apical domain. (**F**) Bar chart showing comparisons of the time difference in polarization between cells with or without overexpression of Tfap2c, Tead4 and RhoA-Q63L in the same embryo, or in control embryos. Scale bars, 15µm.

We have previously shown that apical domain formation requires activation of actomyosin by PKC-Rho GTPase signaling at the 8-cell stage although actomyosin activation alone is insufficient to trigger apical domain formation (Fig. 3B-C, Movie S5) (Zhu et al., 2017). We therefore hypothesized that activation of Rho GTPase might be required concomitantly with the Tfap2c and Tead4 to achieve complete cell polarization. To test this hypothesis, we expressed Tfap2c and Tead4 at the 2-cell stage (together with the Ezrin-RFP as a live apical marker) and constitutively active RhoA-Q63L at the 4-cell stage (Fig. 3A). Strikingly, when all three factors were expressed, complete cap-like apical domains became established at the 4-cell stage (Fig. 3B-C; Movie S6). These prematurely induced apical domains were enriched with Ezrin and Pard6 and thus strongly resembled the apical domains that form normally at the late 8-cell stage (Fig. 3B; Fig. S8B-C). To our knowledge, this is the first time that any premature cell polarization has been reported in the mouse embryo, where this process has been previously considered invariant.

To further confirm these results, we overexpressed all factors in half of the Ezrin-RFP labelled embryos, using the remaining cells as controls (Fig. 3D). In line with our findings from the expression of these factors in the entire embryo, we observed that individual blastomeres targeted with overexpression of the three factors polarized significantly earlier than control blastomeres from the same embryo (Fig. 3E-F; Movie S7-8). We did not observe any difference in the timing of cell division between blastomeres suggesting that polarization at the 4-cell stage by Tfap2c, Tead4 and active RhoA expression is not caused by a delay to cytokinesis (Fig. S8A).

Together our results indicate that the induction of the transcriptional program triggered by Tfap2c and Tead4 alongside the activation of actomyosin downstream of Rho GTPase signaling constitutes the timing mechanism that triggers apical domain formation at a specific stage of preimplantation development.

### Advancing expression of Tfap2c, Tead4 and Rho GTPase advances morphogenesis and cell differentiation

During normal development, polarization at the 8-cell stage is followed by a zippering process in which adjacent apical domains expand and seal their boundaries at the late 16-cell stage, a process essential for blastocyst formation (Zenker et al., 2018). We therefore wished to determine whether the premature cell polarization induced by advancing the onset of Tfap2c/Tead4/RhoA-Q63L expression could also advance the zippering process. To this end, we again induced expression of Tfap2c/Tead4/RhoA-Q63L, to trigger the formation of apical domains at the 4-cell stage, and then followed subsequent development by time-lapse microscopy. The premature formation of the apical domains resulted in premature zippering at the 8-cell stage (Fig. S8B-D). These prematurely zippered sites were enriched with the tight junction protein ZO-1 just as in normal development at the late 16-cell stage (Fig. S8C). These results confirm that advancing the expression of Tfap2c/Tead4/RhoA-Q63L is sufficient to induce the formation of a precocious functional apical domain and also show that advancing cell polarization advances the subsequent step of morphogenesis.

Cell polarization in the mouse embryo is followed by cell fate specification, namely differentiation into the TE cells that inherit an apical domain. Thus, we determined whether changing the timing of cell polarization would affect the timing of TE formation. To this end, we induced apical cap formation at the 4-cell stage, by overexpression of Tfap2c, Tead4 and RhoA-Q63L, and examined the expression of cell differentiation markers. The induction of premature cell polarization induced premature expression of the key TE differentiation transcription factors, Cdx2 and Gata3 (Fig. S8E-H; Movie S9-10). These results suggest that the combined activities of Tfap2c, Tead4 and RhoA-Q63L are sufficient to advance the timing of not only cell polarization but also the cell differentiation program.

### Apical protein centralization is achieved by apical protein clustering

We next wished to define the relative roles of actomyosin activation, Tfap2c, and Tead4 in driving apical domain assembly. The formation of the apical domain does not require canonical symmetry breaking cues but is highly responsive to the dynamics of actin cytoskeleton (Fleming et al., 1986; Johnson, 2009; Korotkevich et al., 2017). As it has been shown that disrupting the actin network abolishes apical domain formation (Sun et al., 2013; Zhu et al., 2017), we interrogated interactions between actin and the apical proteins examined by filming the development of LifeAct-GFP and Ezrin-RFP expressing embryos. Our time-lapse movies revealed that apical domain formation occurred in two steps (Fig. 4A). In the first, centralization step, apical proteins became concentrated around the center of the cell-contact free surface concomitant with local exclusion of actin (Fig. 4A-B). In the second, expansion step, apical proteins accumulated and then expanded to form an apical patch overlaying the actin meshwork before being concentrated in a surrounding ring-like structure (Fig. 4A,C; Movie S11). The initial apical protein centralization step failed to take place in Tfap2c and Tead4 depleted embryos, suggesting that these two transcription factors regulate this first step of apical domain formation (Fig. 2G; Fig. 4D).

**Figure 4.**
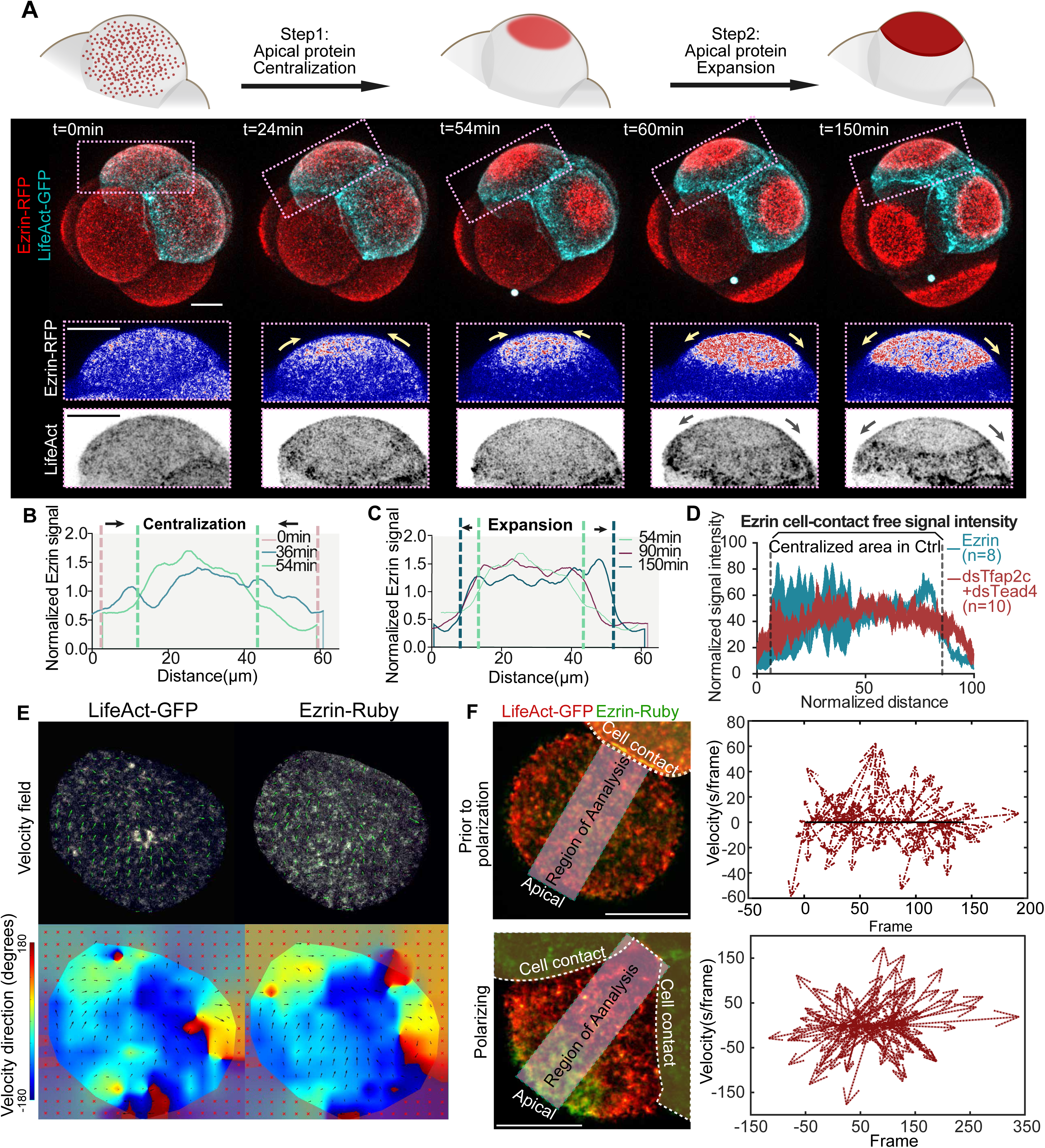
Tfap2c and Tead4 regulate apical domain centralization. **(A)** LifeAct-GFP and Ezrin-RFP dynamics during mid to late 8-cell stage development. Squares denote the magnified regions; yellow arrows show the direction of apical protein movement; grey arrows show the directions of actin ring movements. The apical domain forms in two steps: Step 1: apical polarity proteins become concentrated in the center of the cell-contact free surface. Step 2: the apical proteins continue to accumulate to form an apical domain. Time-lapse represents N=7 total embryos examined. N=4 independent experiments. (**B**) Ezrin-RFP signal at the cell-contact free surface during the apical centralization step in the time-lapse movie shown in **A**. (**C**) Ezrin-RFP signal at the cell-contact free surface during the apical expansion step. For each time-point, Ezrin signal is normalized against the average signal intensity across all measurements, signal extracted from time-lapse in **A**. (**D**) Ezrin-RFP signal at the cell-contact free surface after Ezrin-RFP with dsGFP (as a control, Ctrl) or Tfap2c+Tead4 dsRNAs injections. N=8 cells from N=8 embryos for dsGFP and N=10 cells from N=10 embryos Tfap2c+Tead4 dsRNAs injected cells plotted. N=2 experiments. (**E**) PIV analysis of LifeAct-GFP and Ezrin-Ruby membrane particle movements in the compacted 8-cell stage cells. The directions of the flow for LifeAct-GFP and Ezrin-RFP are coordinated. (**F**) Feather plot shows the time-dependent changes of the sum of vectors in a selected rectanglular area (shown in images on the left) projecting from the cell-cell contact to the cell-contact free surface. Direction of the arrows indicate the direction of the sum of the vectors, and the length of the arrows indicates the velocity of the sum of the vectors. Scale bars, 15µm.

To our surprise, we did not observe obvious actomyosin movements towards the center of cell-contact free surface that could drive apical protein centralization (Movie S11-12). To confirm this observation, we performed time-lapse imaging using high temporal resolution (4s per frame) and analyzed the movements of actin and Ezrin using embryos expressing LifeAct-GFP and Ezrin-RFP. We observed that the actin formed a circumferential, turbulent cortical flow, and that the movement of actin coordinated with the movement of Ezrin (Fig. 4E; Movie S13). However, the overall direction of these flows was not toward the center of the cell-contact free surface (Fig. 4F; Movie S14). Moreover, the inhibition of actomyosin contractility by blebbistatin treatment, impaired such cortical movements but failed to prevent apical protein centralization (Fig. S9A-B; Movie S15)(Maitre et al., 2015; Zhu et al., 2017). These findings suggest that the turbulent actin flow we observed on the cell surface is unlikely to cause apical protein centralization.

Besides the cortical actin flow, we observed that Ezrin forms clusters on the cell-contact free surface, and that the centralization of Ezrin is accompanied by an exponential growth of the size of Ezrin clusters in the center of the cell-contact free surface (Fig 5A-C; Movie S12). This result suggests the possibility that the centralization of apical proteins is achieved by the conjugation of apical protein clusters on the cell-contact free domain. We also observed a phase-dependent correlation of Ezrin and actin dynamics during the period of apical domain centralization. Initially, when the Ezrin cluster was small, actin and Ezrin localization exhibited a moderate positive correlation (Fig. 5D). Live imaging revealed that the conjugation of Ezrin clusters happened during the merging and splitting of actin clusters that was not prevented by blebbistatin treatment (Fig. 5E), suggesting that actin turnover and consequent actin cytoskeleton remodeling may trigger apical protein clustering and centralization. To test this idea, we treated mid 8-cell stage embryos with two actin inhibitors, firstly Jasp(Bubb et al., 1994), a drug that prevents actin de-polymerization; and secondly CK666(Sun et al., 2013), an inhibitor that prevents Arp2/3 mediated actin polymerization. In support of our idea, Jasp and CK666 treatment consistently blocked the growth of Ezrin clusters and therefore the centralization of Ezrin proteins (Fig. 5F-G; Fig. S9C-D). Interestingly, overexpression of RhoA-Q63L reduced the formation of clusters but increased membrane localization of Ezrin (Fig. S9E-F); on the contrary, depletion of RhoA resulted in ectopic clustering of actin and Ezrin on the cell membrane (Fig. S9G-H). These data suggest that RhoA signaling remodels the cortical property that negatively regulates cluster formation but prompts apical protein membrane localization. In addition, we also observed a correlation between cell curvature and the apical protein clustering site (Fig. S9I), which suggests that apical protein clustering preferentially happen at place with high curvature. To test this idea, we elongated early 8-cell stage cells to induce the asymmetry of cell curvature, using a method we have previously used (Methods)(Gray et al., 2004), followed by the tracking of the position of the apical domain using the time-lapse imaging. We found that the apical domain preferentially develops on the poles with high curvature (Fig. S9J). These results suggest that apical clustering responds to membrane geometry leading it to develop on the cell-contact free surface (Korotkevich et al., 2017).

**Figure 5.**
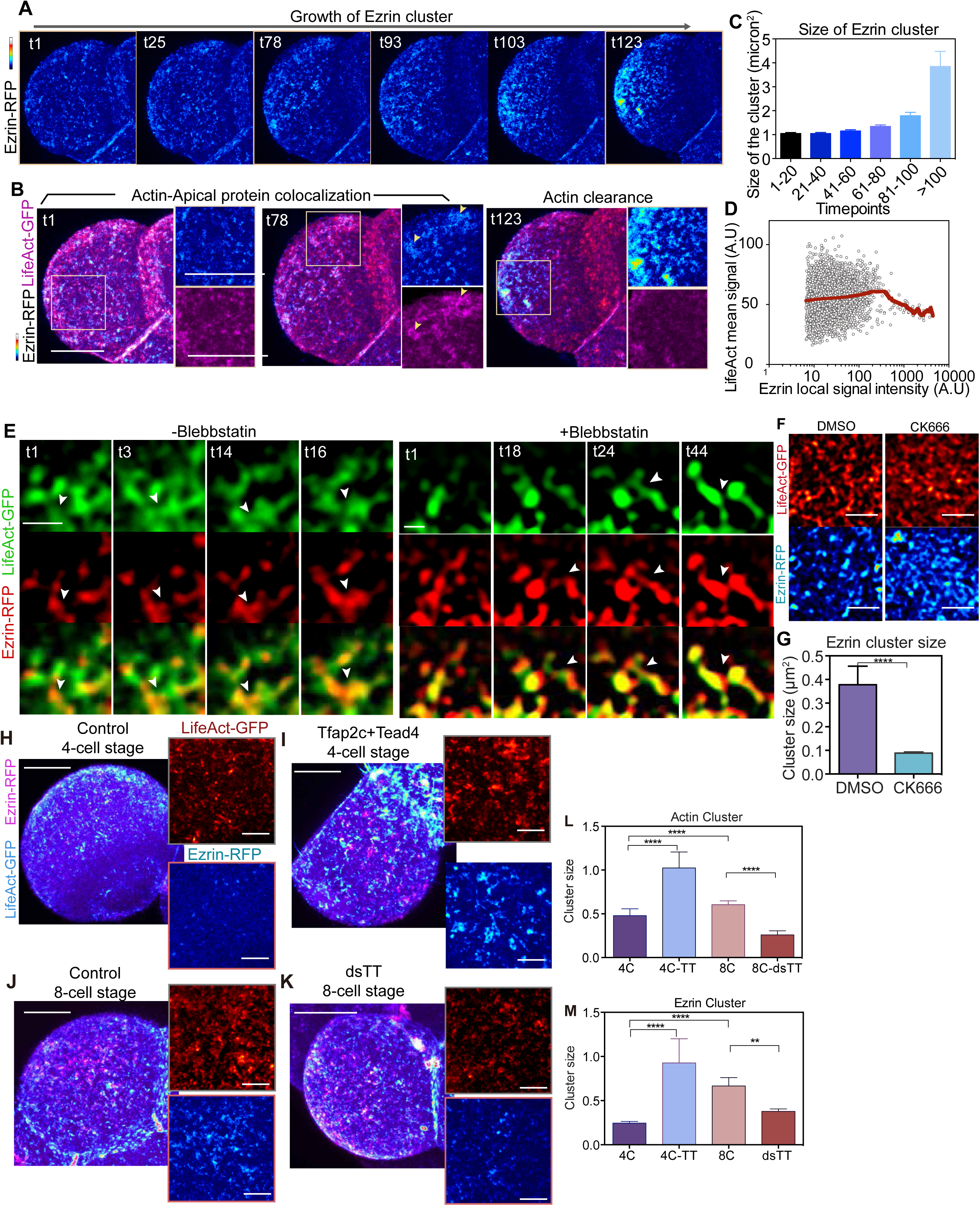
Clustering of apical proteins drives the centralization of the apical domain. **(A,B)** Snapshots from a time-lapse movie monitoring the dynamics of Ezrin-RFP and LifeAct-GFP. Squares indicate the magnified region shown on the right of each image. Note that the Ezrin clusters become larger with time. Yellow arrows indicate the positive correlation of the localization between actin and Ezrin clusters. Quantification is shown in **C**. (**C**) Quantification of the size of Ezrin clusters during cell polarization. For each dataset, more than 1500 clusters were analyzed. Data present as a bar chart showing mean ± S.D. N=2 independent experiments. (**D**) Quantification of mean LifeAct-GFP signal intensity against the total Ezrin signal intensity in a Ezrin cluster (described as “Ezrin local signal intensity”). Total Ezrin signal intensity was calculated by multiplying the mean Ezrin-RFP signal intensity with the size of the Ezrin cluster. The LOWESS curve (in red) was calculated to show the approximate correlation between Ezrin-RFP and LifeAct-GFP. Overall the relationship shows two phases: when Ezrin level is low, a slight positive correlation can be seen with LifeAct, but when Ezrin level is high (with a threshold of 366.0 A.U⋅μm^2^), LifeAct shows a negative correlation with Ezrin. Each dot represents one Ezrin cluster. N=2 independent experiments. (**E**) Snapshots from a time-lapse movie showing LifeAct-GFP and Ezrin-RFP localization with 3s time interval with or without Blebblstatin treatment. Arrows indicate the merging of adjacent Ezrin clusters as a result of actin polymerization. Scale bar, 1μm. Images represent N=5 regions examined from N=5 cells for each condition. N=3 independent experiments. Blebbistatin treatment did not prevent the clustering of actin or Ezrin proteins. (**F**) Localization of LifeAct-GFP and Ezrin-RFP in embryos treated with DMSO (control) and CK666. Scale bars, 5μm. Quantifications are shown in **G**. (**G**) Quantifications of actin cluster size in cells treated with DMSO and CK666. (**H**-**K**) Localization of LifeAct-GFP and Ezrin-RFP in embryos injected with LifeAct-GFP and Ezrin-RFP mRNA, with or without Tfap2c and Tead4 mRNA at the late 4-cell stage, or embryos injected LifeAct-GFP and Ezrin-RFP mRNA with or without dsRNA targeting Tfap2c and Tead4 at the 8-cell stage. Magnified regions are shown on the right. Quantifications are shown in **L** and **M**. (**L**,**M**) Quantifications of the size of actin (**L**) or Ezrin (**M**) clusters in embryos shown in (**H-K**). Data shown as mean ± S.E.M. ***p<0.001, ****p<0.0001, One-way ANOVA test. N=3 4-cell stage control cells; N=4 Tfap2c+Tead4 mRNA overexpressed cells; N=4 control 8-cell stage cells; N=3 dsTfap2c+dsTead4 injected cells. Scale bars for images on the left in **H-K**, 15µm; for magnified images on the right, 5µm.

Together, these observations indicate that apical protein centralization is mediated by the clustering of apical proteins that is regulated by both actin turnover and myosin activity.

### Tfap2c and Tead4 control apical protein clustering

We next wished to determine the mechanisms by which Tfap2c and Tead4 regulate apical domain centralization. To this end, we examined how the level of Tfap2c-Tead4 activity would influence the configuration of actin and Ezrin on the apical surface. We found that depletion of Tfap2c and Tead4 resulted in smaller actin and Ezrin cluster sizes by the mid 8-cell stage. Conversely, overexpression of Tfap2c and Tead4 significantly increased the actin and Ezrin cluster sizes at the late 4-cell stage (Fig. 5H-M). These results suggest that Tfap2c and Tead4 regulate apical domain centralization by influencing expression of proteins that modulate apical protein clustering.

To gain further mechanistic insight into how Tfap2c-Tead4 regulate apical domain formation, we carried out RNA-sequencing of early 8-cell stage embryos depleted of Tfap2c and/or Tead4. For each group of embryos (control *GFP* RNAi, *Tfap2c* RNAi, *Tead4* RNAi, *Tfap2c*/*Tead4* co-RNAi), we collected two biological replicates with 10 embryos per sample, using embryos of two strains to eliminate any effect of the genetic background (Fig. 6A, Methods). The effect of different treatments on the global gene expression pattern was highly reproducible between biological replicates and between genetic backgrounds (Fig. S10A). Differential gene expression analysis (with a 2-fold cut-off) showed that depletion of Tfap2c led to the downregulation of 749 or 929 genes depending on the strain, whereas depletion of Tead4 led to downregulation of 242 or 314 genes (Fig. S10B-C). The co-depletion of Tfap2c and Tead4 led to an additional 135 or 95 genes being downregulated compared to single knockdown embryos, depending on the strain (Fig. S10B-C). Among the genes consistently downregulated in double-knockdown embryos of both strains, a significant proportion was associated with actin polymerization. These include the Arp2/3 complex, the tropomyosin complex, Marcks proteins, and the FREM family member Ebp4.1l5, and Cdc42 effector protein family members (Borg)(Vong et al., 2010) (Fig. 6B; Table S3). The levels of expression of these factors are upregulated between the 2- to 8-cell stages and correlate with the growth of Ezrin clusters over this period. Importantly, the depletion of Tfap2c and Tead4 eliminates their expression resulting in the reduced apical cluster size (Fig. 6B; Fig. 5H-M). To further validate whether the expression of these downstream targets are responsible for apical domain formation, we depleted two of the highly expressed candidates, Arpc1b and Marcksl1, and found that this led to the impaired apical domain formation (Fig. 6C-E). These results together suggest that Tfap2c and Tead4 regulate apical protein clustering by regulating the actin network, to centralize the apical proteins.

**Figure 6.**
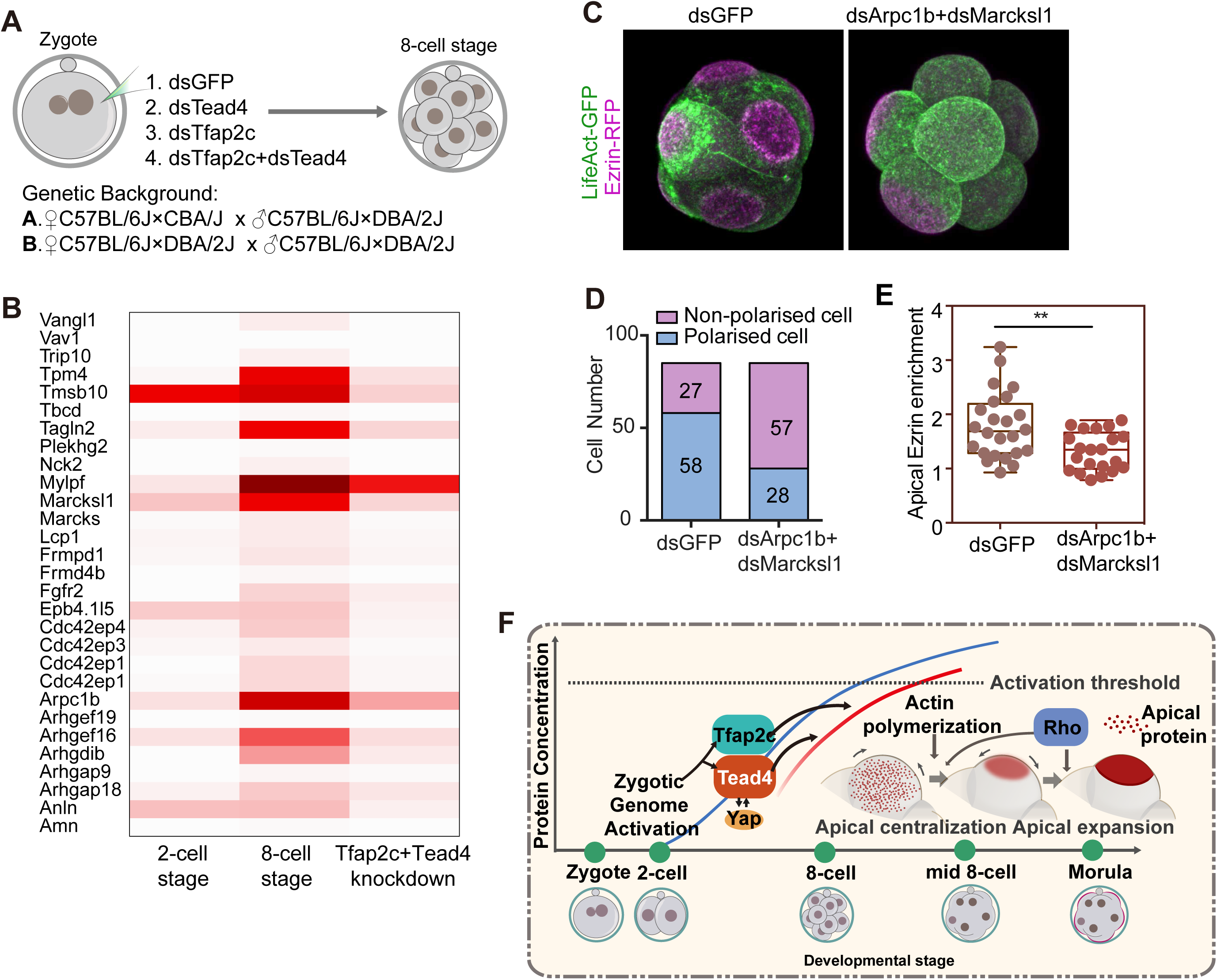
Tfap2c and Tead4 regulate the expression of actin regulators. (**A**) Schematic view of the experimental conditions and genetic backgrounds used for RNA-sequencing. (**B**) Heatmap showing the expression profile of selective cytoskeleton regulators downstream of Tfap2c and Tead4. (**C**) Representative images of embryos injected with dsGFP or dsRNA targeting Arpc1b and Marcksl1 and examined at the late 8-cell stage to reveal the localization of LifeAct-GFP and Ezrin-RFP. (**D**) Quantification of the number of cells that with or without the apical domain at the late 8-cell stage in dsGFP and dsArpc1b+dsMarcksl1 groups. Number in each bar represents the number of embryos in each category. (**E**) Quantification of apical enrichment of Ezrin-RFP in cells injected with dsGFP or dsArpc1b+dsMarckl1. Each dot represents an analyzed cell. (**F**) Summary of the results. Taken together our results suggest a model of how *de novo* cell polarization in the mouse embryo is controlled through Tfap2c, Tead4 and RhoA activity. Tfap2c and Tead4 proteins gradually accumulate from the 2-cell to the 8-cell stage. Tead4 expression induces nuclear localization of Yap generating the Tead4-Yap transcription complex. Tfap2c and Tead4-Yap induce expression of essential regulators and drive the central enrichment of apical protein components. Then Rho protein activity remodels the properties of the actin cytoskeleton leading to the expansion of the apical domain, triggering the formation of the final apical cap and thereby inducing cell polarization.

In summary, our results show that the activation of actomyosin by Rho GTPases concurrent with the transcriptional activity of Tfap2c and Tead4 triggers cell polarization and the segregation of the TE and ICM fates. Together, these three factors control the initiation of expression of transcription factors required for TE formation and the establishment of the apical domain to execute the first cell fate decision (Fig. 6F).

## Discussion

The first bifurcation of cell fate in the mammalian embryo is a fundamental step that separates progenitors of the embryonic and extra-embryonic tissues. The establishment of the apical domain is the primary trigger for this process (Bedzhov et al., 2014; Chazaud and Yamanaka, 2016; Johnson and Ziomek, 1981a; Korotkevich et al., 2017). Although great progress has been made in understanding the connections between cell polarity and cell fate (Anani et al., 2014; Hirate et al., 2013; Samarage et al., 2015), little is known about how the establishment of the apical domain itself is triggered with such temporal specificity during development. Here, we provide evidence denoting the importance of zygotic genome activation in establishing the timing of cell polarization and demonstrate that the zygotic expression of the transcription factors Tfap2c and Tead4 is necessary for this process. We further identified a functional redundancy in their regulation of cell polarization, which can explain why this role was not discovered in previous studies where single knockout embryos were examined (Nishioka et al., 2008; Winger et al., 2006; Yagi et al., 2007). Our finding that the abundance of Tfap2c and Tead4 proteins increases after zygotic genome activation and correlates with the exposure of their DNA binding sites to open chromatin regions (Fig.S4) (Wu et al., 2016) indicates that their zygotic expression accounts for their functional activity. Accordingly, we show that inducing upregulation of Tfap2c and Tead4 in conjunction with Rho GTPase-mediated activation of actomyosin is sufficient to establish the apical domain and this allowed us, for the first time, to advance the timing of cell polarization. In turn, this advances the expression of downstream lineage-specific transcription factors that establish TE identity. This supports the idea of feedback loops between cellular events and the robust segregation of the first lineages (Cao et al., 2015; Yagi et al., 2007).

Apico-basal cell polarization at the 8-cell stage of the mouse embryo has often been viewed as a model for epithelial polarization. However, the mechanism establishing the apical domain is quite distinct from many other cell types as it can be formed in the absence of external cues such as those provided by the extra-cellular matrix or through cell adhesion (Korotkevich et al., 2017). Such spontaneous symmetry breaking properties have also been deployed by a broad array of tumor cells, but their underlying mechanisms remain elusive (Lorentzen et al., 2018). Here we show that the apical proteins form clusters at the cell-contact free surface and the conjugation of apical protein clusters, controlled by Tfap2c and Tead4 activity, leads to the apical protein polarization. Actomyosin flow has been reported to trigger cortical polarization in the C.elegans zygote. We show that although actomyosin flow helps to restore the apical domain after cytokinesis, at the 16-cell stage, we did not observe the actomyosin flow to contribute to the establishment of cell polarity at the 8-cell stage. Such mechanistic differences may correlate with the distinct time-scales of cell polarity establishment between different species, and the utilization of actomyosin flow may account for the more rapid establishment of a cortical domain in the worm embryo and in recovery of the domain in the 16-cell stage mouse embryo.

The clustering of apical proteins could be achieved through a change in the binding kinetics of apical proteins to lipids and actin. Indeed, we observed a strong positive correlation between Ezrin/Pard6 and PIP2 localization throughout the apical domain formation process (data not shown). The biophysical properties of the actin network, determined by actin turnover and myosin activity (mediated by Rho GTPases), may provide the driving force directing lipid-apical protein clustering. It has been reported that the PIP2 lipid prefers to cluster in areas with high cell curvature (Lin et al., 2018). Accordingly we also found a positive correlation between cell curvature and the position of the apical domain. These results may account for the centralized positioning of the apical domain on the cell contact-free surface. PIP2 regulated apical protein localization may provide a mechano-sensing mechanism independent of adherens junction to establish the apical domain, as observed in E-cadherin knockout cells (Korotkevich et al., 2017), it is of future interest to examine this possibility.

In summary, our results provide mechanistic insight into the timing and establishment of de novo cell polarization in the mouse embryo, the critical event for the transition from totipotency to pluripotency.

## Acknowledgement

We are grateful to David Glover and Marta Shahbazi for valuable comments on the manuscript; Ed Munro for helpful discussion of the project; Stavros Malas for providing the Gata3-GFP transgenic line. This work was supported by Wellcome Trust (098287/Z/12/Z), ERC (669198) and Leverhulme Trust (RPG-2018-085) grants to M.Z.G.

## Author contributions

Conceptualization: M.Z and M.Z.G.; Investigation: M.Z., P.W., C.H; Writing: M.Z and M.Z.G.; Supervision: M.Z.G., N.J.

## Animals

This research has been carried out following regulations of the Animals (Scientific Procedures) Act 1986 - Amendment Regulations 2012 - reviewed by the University of Cambridge Animal Welfare and Ethical Review Body. Embryos were collected from F1 females (C57BI6xCBA) that had been super-ovulated by injection of 7.5 IU of pregnant mares’ serum gonadotropin followed by human chorionic gonadotropin (Intervet) 48 h later. F1 females were mated with F1 males.

## Mouse embryo culture and inhibitor treatments

Embryos were recovered at the zygote or 2-cell stage in M2 medium and subsequently transferred to KSOM medium for long-term culture, as described previously (Zhu et al., 2017).

Inhibitor treatment: Puromycin (Invivogen, ant-pr-1) was diluted in KSOM to a working concentration of 10μg/ml. Cycloheximide (Sigma-Aldrich, C7698) was dissolved in DMSO and diluted in KSOM to a working concentration of 20μg/ml. 5,6-Dichlorobenzimidazole 1-b-D-ribofuranoside (DRB; Sigma-Aldrich, D1916) and Triptolide (Cayman Chemical, CAY11973) were dissolved in DMSO and diluted in KSOM to a working concentration of 50μM (DRB) or 5μM (Triptolide). C3-transferase was dissolved in distilled water and diluted in KSOM to 7μg/μl. For the control groups, the same dilutions of the vectors of different inhibitors were added to the medium.

## Blastomere resection

The resection procedure was performed as previously described(Zernicka-Goetz, 1998). Briefly, the zona pellucida was removed for both 2-cell stage and 4-cell stage embryos. Embryos were transferred to a 1% agarose coated petri-dish covered by M2-medium containing 2µM Cytochalasin D (Sigma-Aldrich, C8273) prior to the resection procedures. For the 2-cell embryo, the embryo was first elongated by using a thin glass capillary with a flame-polished end. One of the blastomeres was resected using a thin glass needle, leaving approximately 30-40% of the cytoplasm (cytoplast) attached to its sister cell (Fig. 2B). Cell volume measurement was performed by applying a 3D polygon ROI around the periphery of the structure (indicated by Ezrin-RFP) throughout the Z-stack. The measurement was performed by using Icy software. For the 4-cell stage resection, the 4 blastomeres were first transferred to Calcium-Magnesium free M2 medium for 5 min and cells were dissociated by pipetting, as previously described (Graham et al., 2014). All four blastomeres were elongated using a thin glass capillary, and only two were resected (Fig. 2C). The small and control cells were transferred to M2 medium immediately after resection. The whole resection process for each embryo took up to 5 min with a survival rate greater than 80%. The small and control cells were transferred to KSOM medium for long-term culture.

## Blastomere elongation

Elongation of 8-cell stage blastomeres were performed using the method described previously(Gray et al., 2004), the cells from an early 8-cell stage embryo (0-1hr post cell division) were disassociated as described in blastomere resection experiments. The single cell was placed in KSOM supplemented with containing sodium alginate (0.5%). The cell was then elongated by pipetting with a thin glass capillary. Immediate after the elongation procedure, a few drops of a 1.5% CaCl2 (0.3g CaCl2 dissolved in 20.0 ml 0.15M NaCl) were added, leading to the gelling of KSOM surrounding the cell and hence the maintenance of cell shape. The excessive CaCl2 was removed by replacing the medium with KSOM.

## Microinjection

Microinjection was carried out as described previously (Zernicka-Goetz et al., 1997). In brief, embryos were placed in M2 medium on a glass slide with a depression and covered by a drop of mineral oil. Microinjection was performed with an Eppendorf Femtojet Microinjector. Negative capacitance was used to facilitate penetration through the membrane. dsRNA was injected at a concentration of 1μg/μl. Synthetic mRNAs were injected at the following concentration: Ezrin-Ruby (400ng/μl); Ezrin-Venus (400ng/μl); Tfap2c (15ng/μl); Tead4 (15ng/μl); RhoA-Q63L (3ng/μl); GFP-Myl12b (300ng/μl); Cas9 (100ng/μl). All sgRNAs were injected at 25ng/μl.

## Preparation of DNA Constructs

pRN3P was used as the vector for all constructs as previously described (Zernicka-Goetz et al., 1997). To construct pRN3p-Tead4, Tead4 was amplified from mouse kidney cDNA and cloned into the pRN3p vector. To construct pRN3p-Tfap2c, Tfap2c cDNA was purchased from Origene (MR207174) and cloned into the pRN3p vector. pRN3p-Cas9 was a gift of J. Na (Tsinghua University, School of Medicine). Ezrin-Ruby, Ezrin-Venus, LifeAct-Ruby, GFP-Myl12b, RhoA-Q63L were as previously described (Zhu et al., 2017). All primers for constructs preparation are listed in Table S4.

## mRNA, dsRNA, sgRNA preparation

For mRNA preparation, constructs for each mRNA were linearized using a restriction site downstream of the poly-A region. In vitro transcription was performed using the mMessage mMachine T3 kit (Thermo Fisher, AM1348) following the manufacturer’s instructions. mRNAs were purified using the lithium chloride precipitation method. For sgRNA preparation, the sequences of sgRNAs were designed using CRISPR design tool website (http://cirpsr.mit.edu). The DNA fragment containing T7 promoter, crRNA and sgRNA sequence were amplified using the Geneart gRNA kit (Thermo Fisher, A29377). sgRNAs were in vitro transcribed and purified using the gRNA Clean Up Kit (Thermo Fisher, A29377), following manufacture’s instructions.

All dsRNAs were designed using the E-RNAi website (Horn and Boutros, 2010) and were 350-500bp in length. The specific targeting regions for each dsRNA were amplified from a mixture of mouse kidney, lung, liver cDNAs. The in vitro transcription reactions were performed using the MEGAscript T7 transcription kit (Thermo Fisher, AM1334) following the manufacturer’s instructions. dsRNAs were purified by lithium chloride precipitation. All primers for dsRNA preparation are listed in Table S3.

## Immunofluorescence

Embryos were fixed in 4% PFA at room temperature for 20min, and washed in PBST (0.1% Tween in PBS) three times. The embryos were then permeabilised in 0.5% Triton X-100 in PBS for 20 min at room temperature, washed in PBST three times, transferred to blocking solution (3% bovine serum albumin) for 2h and incubated with primary antibodies (diluted in blocking solution) at 4 °C overnight. After the incubation, embryos were washed in PBST and incubated with secondary antibodies (1:500 in blocking solution) for 1h at room temperature. Embryos were stained with DAPI (1:1000 dilution, in PBST, Life Technologies, D3571) for 15 min, followed by two washes in PBST. Primary antibodies: rabbit polyclonal anti-Pard6b (Santa Cruz, sc-67393, 1:200); mouse monoclonal anti-GFP (Nacalai Tesque Inc., 04404-84, 1:500). mouse monoclonal anti-Tfap2c (Santa Cruz, sc-12762, 1:200); goat monoclonal anti-Tfap2c (R&D Systems, AF5059-SP, 1:200); rabbit monoclonal anti-Tead4 (Abcam, ab97460, 1:200); mouse monoclonal anti-Tead4 (Abcam, ab58310, 1:100); goat monoclonal anti Sox17 (R&D Systems, af1924); mouse monoclonal anti Cdx2 (Launch Diagnostics, MU392-UC (Biogenex), 1:200); rabbit monoclonal anti Nanog (Abcam, ab80892, 1:200); mouse monoclonal anti-Tjp1 (Thermo Fisher Scientific, 33-9100, 1:200); rabbit monoclonal anti-phosphorylated-Yap (Cell Signaling Technologies, 4911S, 1:200); mouse monoclonal anti-Yap (Santa Cruz, sc-101199, 1:200); rabbit monoclonal anti-di-phosphorylated MRLC (Cell Signaling Technologies, 3674P, 1:100); goat polyclonal anti-Amot (Santa Cruz, sc-82491, 1:1000). Secondary antibodies: Alexa Fluor 568 Donkey anti-Goat (A-11057, ThermoFisher Scientific); Alexa Fluor 488 Donkey anti-Mouse, (A-21202, ThermoFisher Scientific); Alexa Fluor 568 Donkey anti-Mouse (A10037, ThermoFisher Scientific); Alexa Fluor 647 Donkey anti-Mouse (A31571, ThermoFisher Scientific); Alexa Fluor 568 Donkey anti-Rabbit (A10042, ThermoFisher Scientific); Alexa Fluor 647 Donkey anti-Rabbit (A-31573, ThermoFisher Scientific).

## Real-time PCR

RNA was extracted from 8-cell stage embryos using the Arcturus PicoPure RNA isolation kit (Arcturus Bioscience). RT-PCR was performed using a StepOne Plus Real-time PCR machine (Applied Biosystem). The expression level was calculated using ddCT methods, normalised to a Gapdh PCR reaction and the endogenous control group. The primers used for RT-PCR are listed in Table S4.

## Imaging and data processing

Imaging was carried out on a Leica-SP5 or a Leica-SP8 confocal using a Leica 1.4 NA 63X oil (HC PL APO) objective. Images were processed with Fiji software (Schindelin et al., 2012). For the analysis of nucleo-cytoplasmic signal intensity ratio, the region of the nucleus, and a cytoplasmic region of the same size, were cropped and the mean signal extracted using the Fiji ROI function. To normalise signals to the level of DAPI fluorescence, the Fiji ROI function was used to extract the nuclear region stained to reveal specific proteins and for the equivalent DAPI channel and normalised using the formula: *I*(protein of interest)/(DAPI). For Ezrin apical enrichment analysis, a freehand line of the width of 0.5μm was drawn along the cell-contact free surface (apical domain), or cell-contact (basal) area of the cell, signal intensity was obtained via ROI function of Fiji. The apical/basal signal intensity ratio is calculated as: *I*(apical)/*I*(basal). Cells on the same plane were subjected to this analysis. Compaction was assessed by measuring the inter-cellular blastomere angle in the mid-plane between adjacent cells (as described previously(Zhu et al., 2017)) by using the Fiji angle function.

For live-imaging, time-lapse recordings of embryos were carried out using a spinning disk or a Leica-SP5 scanning confocal. For the blastomere resection experiment, time-lapse frames were acquired every 1hr; for other live-imaging experiments, time-lapse frames were acquired every 20-30min. Images were acquired using a 3-4μm *Z*-step. Images were processed with Fiji software. Correlations were calculated using Prism software (http://www.graphpad.com).

## Particle Image Velocimetry (PIV) analysis

PIV analysis was performed using PIVlab MATLAB algorithm (pivlab.blogspot.de). The sequential images with 3-4s/frame were used as input. 2-pass analysis with 80×40 pixel was used, and the images were analyzed with A-B,B-C,.. sequence. A mask demarcating the edge of the cell were applied to the image before processing the analysis.

## Statistics

Statistical methods are indicated for every experiment in the corresponding figure legends. Qualitative data is presented as a contingency table and was analyzed using Fisher’s exact test. Normality of quantitative data was first analyzed using D’Agostino’s K-squared test. A one-sample t-test was used to test whether an observed distribution followed a hypothetical mean. If data showed a normal distribution, then for comparison of two or multiple samples, an unpaired two-tailed Student’s *t* test (two experimental groups) or a One-way ANOVA test (more than two experimental groups) was used to analyze statistical significance. Differences in variances were taken into account by performing a Welch’s correction. For data that did not present a normal distribution, a Mann–Whitney *U*-test (two experimental groups) or a Kruskal–Wallis test with a Dunn’s multiple comparison test (more than two experimental groups) was used to test statistical significance. To determine the influence of different groups in multiple variants, two-way ANOVA was performed. Statistical analyzes were performed using Prism software (http://www.graphpad.com).

## RNA extraction and sequencing

For sample collection, 10 8-cell stage embryos injected with dsRNAs were treated with acidic tyrode solution to remove zona pellucida, washed in PBS (without Ca2+ and Mg2+) and transferred to transferred to hypotonic lysis buffer (Amresco, M334). mRNA was reverse transcribed by SuperScript II and pre-amplified using Smart-seq2 protocol as described previously (Picelli et al., 2014). Pre-amplified cDNA was fragmented by Tn5 enzyme, followed by library generation using TruePrep® DNA Library Prep Kit V2 for illumina Kit (Vazyme, TD501-503). Sequencing was performed on HiSeq X Ten platform.

## RNA-sequencing data Processing

Raw reads with adaptors, low-complexity or low-quality were trimmed by trim_galore. Then clean data were mapped to mouse genome (mm10) by STAR. FPKM (Fragments Per Kilobase per Million mapped reads) of Refseq genes were calculated by Cufflinks. Htseq-count was used to count the mapped reads number. Subsequent reads number was used to perform differential gene expression analysis by using R packages “DESeq2” (Fold change >2, P value < 0.05). Heatmap and volcano plot were graphed using R (http://www.r-project.org/).

---

**Figure S1.**
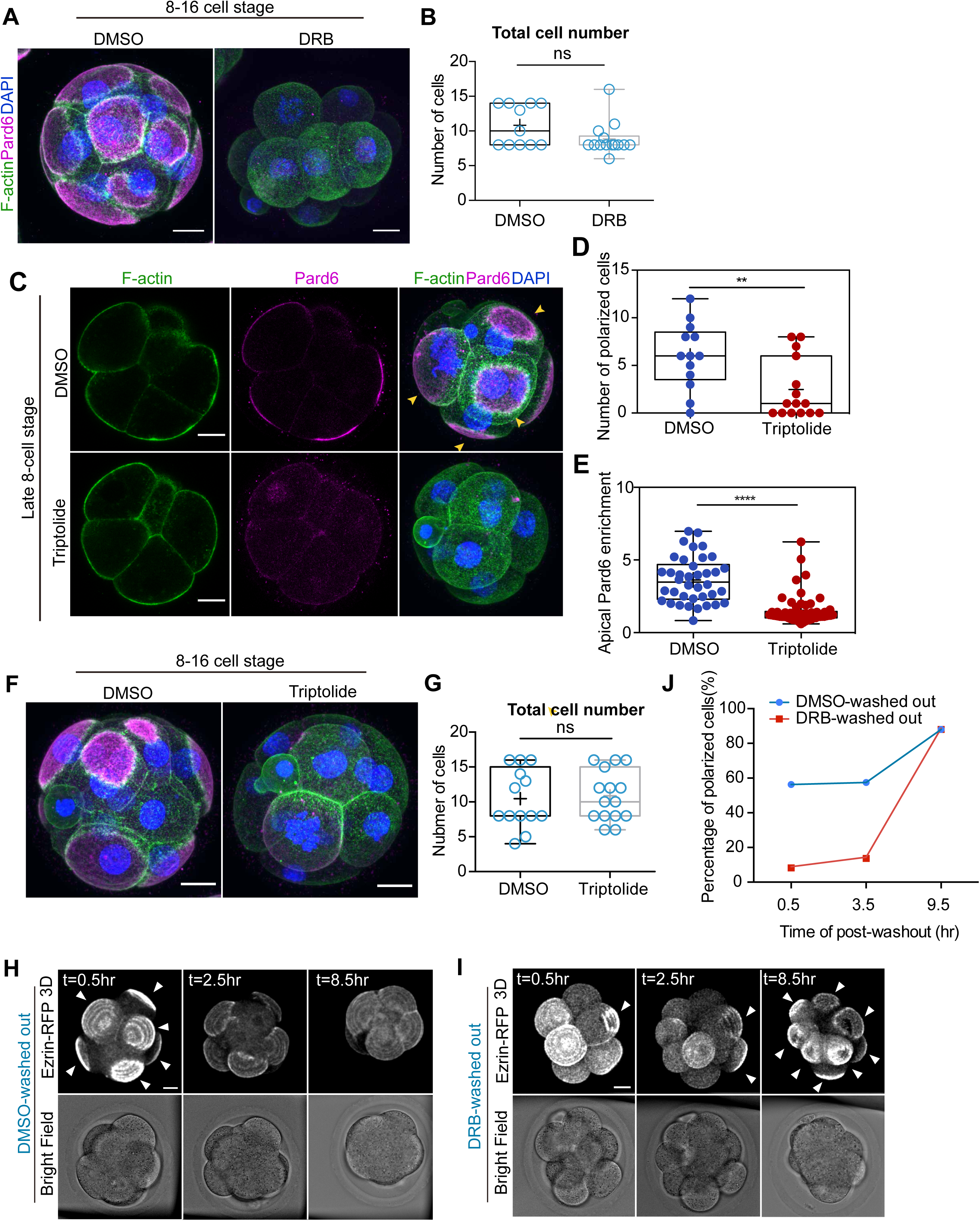
Inhibition of transcription abolished apical domain formation without affecting actomyosin polarization. (**A**) DMSO- or DRB-treated embryos at 8-16 cell stage examined to reveal F-actin, Pard6 and DNA. (**B**) Total cell numbers in embryos treated with DMSO or DRB from 4/8-cell to the late 8-cell stage. Each dot represents an analyzed embryo. ns, not significant; Mann-Whitney test. (**C**) DMSO- or Triptolide-treated embryos at the late 8-cell stage analyzed for F-actin, Pard6 and DNA. (**D**) Quantification of polarized cells in DMSO- and Triptolide-treated embryos at the 8-16 cell stage. **p=0.0088, Student’s t-test. N=13 embryos for DMSO, N=15 embryos for Triptolide. N=2 independent experiments. (**E**) Extent of cell polarization in each cell quantified by the intensity of apical enrichment of Pard6 (Star Methods) in cells treated with DMSO (control) or Triptolide from the early 8-cell stage. Each dot represents an analyzed cell. ****p<0.0001, Mann-Whitney test. (**F**) DMSO- or Triptolide-treated embryos analyzed at 8-16 cell stage for F-actin, Pard6 and DNA. (**G**) Quantification of total cell number in DMSO- and DRB-treated embryos. Each dot represents an analyzed embryo. ns, not significant in Student’s t-test. For **B**, **D**, **E** and **G**, Data are shown as individual data points with Box and Whisker plots (lower: 25%; upper: 75%; line: median; whiskers: min to max). (**H**,**I**) Snapshots from time-lapse movies of embryos after washing out DMSO. (**J**) Line chart shows percentage of polarized cells at different post-wash out time-points in DMSO washed-out (control) or DRB washed-out groups. For t=0.5h, N=87 cells for DMSO and N=97 cells for DRB group were analyzed; for t=3.5h, N=101 cells for DMSO and N=102 cells for DRB group were analyzed; for t=9.5h, N=109 cells for DMSO and N=107 cells for DRB were analyzed. For all analyses, N=11 embryos for DMSO group were analyzed; N=13 embryos for DRB treated group were analyzed. N=2 independent experiments. Scale bars, 15µm.

**Figure S2.**
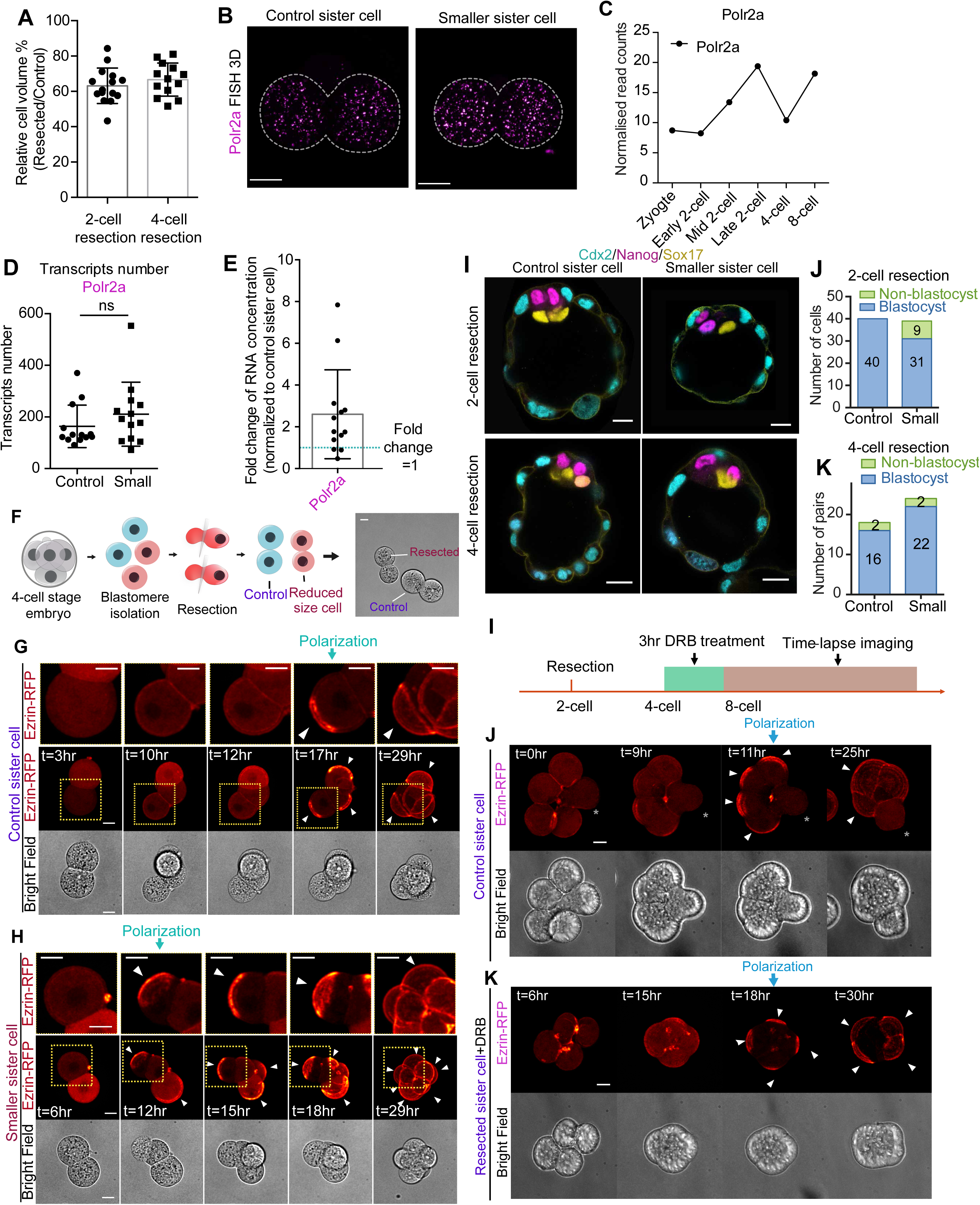
Reduce cell size elevates the concentration of transcripts and induces premature cell polarization. (**A**) Quantification of cell volume in small sister cells/pairs in relation to control sister cells/pairs generated from experiments illustrated in Fig. 1F and Supplementary Fig. S2F. Data is shown as the mean ± S.D.. Each dot represent one analyzed embryo. N=6 independent experiments. (**B**) Control or small sister cells from the same 2-cell stage embryo were cultured for one day and single molecular fluorescence in situ hybridization was used to determine the transcripts for housekeeping gene Polr2a. Quantifications are shown in **C**, **D**, and **E**. (**C**) Transcripts expression profile of Polr2a. Data retrieved from published dataset(Deng et al., 2014). (**D**) Quantification of transcripts number of Polr2a in control and small sister cells (resection performed at the 2-cell stage) at the late 4-cell stage. ns, not significant, Mann-Whitney test. Each dot represents an analyzed embryo. N=13 control or smaller sister cells. N=3 independent experiments. (**E**) Fold change of RNA concentration in small sister cells compared to control sister cells. The RNA concentration is defined as the number of transcripts divided by the cell volume. The RNA concentration of small sister cells divided by the RNA concentration of the control sister cells was used to reveal the fold change. Data is shown as the mean ± S.D. Each dot represents one embryo. (**F**) Schematic diagram of blastomere resection at the 4-cell stage. Cells were separated and two resected; the two small cells and two control cells were then re-aggregated. (**G, H**) Time-lapse movies of control or small sister pairs from experiment in **F**. (**I**) Scheme of pulsed DRB treatment in 2-cell stage resection experiment. Resection performed at the mid 2-cell stage (as indicated in Fig. 1F). Resected cells were subjected to DRB treatment for 3 hr at the late 4-cell stage followed by time-lapse imaging. (**J**, **K**) Time-lapse imaging of control sister cells (**K**) and smaller + DRB-treated sister cells (**J**) in the experiment shown in **I**. Arrows indicate apical domains. Asterisks, cytoplasm from resected cells. Scale bars, 15µm.

**Figure S3.**
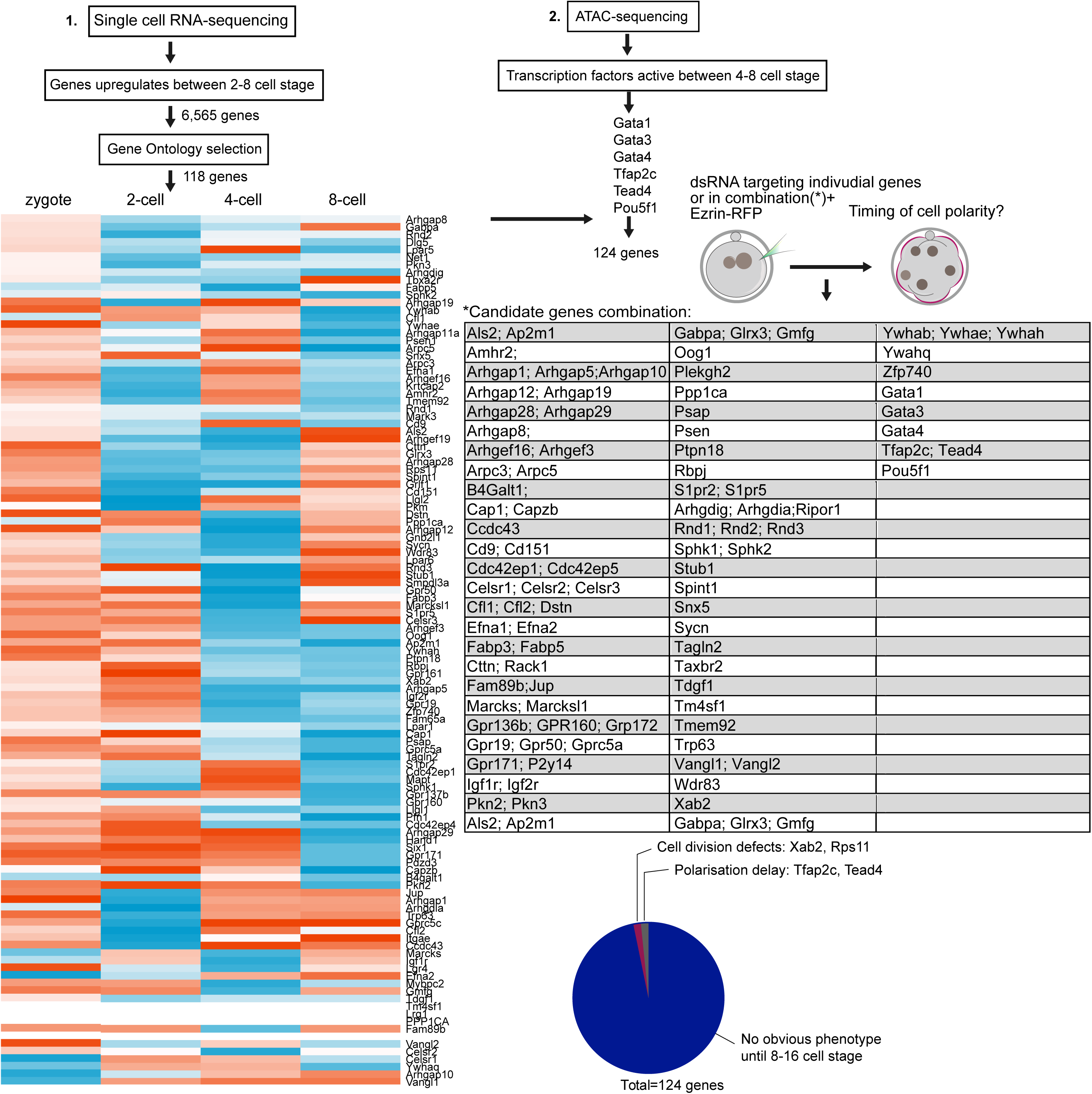
Functional screen to identify regulators of the timing of cell polarization. Candidate regulators were selected based on two hypotheses. The first hypothesis proposes that activators of cell polarization accumulate after zygotic genome activation at the 2-cell stage, to trigger cell polarization. To select candidates based on this hypothesis, single cell RNA-sequencing data was analyzed(Goolam et al., 2016) and transcripts that were upregulated between the 2- to 8-cell stage pooled; their gene ontology was analysed and 118 candidate genes selected falling under the categories of “Rho-GTPases regulators”, “actin cytoskeleton regulators”, and “cell polarity regulators”. The second hypothesis proposes that stage specific transcription factors activate essential regulators of cell polarization around the early 8-cell stage. To select candidates based on this hypothesis, we analyzed ATAC-seq data (Wu et al., 2016) and transcription factors active between the 4-8 cell stages (in total 6 transcription factors were selected). For the 124 combined candidates, one or two dsRNAs targeting individual candidates or combinations were injected at the zygote stage and the timing of the establishment of cell polarization determined by imaging Pard6 at the late 8-cell stage. Depletion of 2 candidates (Xab2, Rps11) lead to cell division defects, and the depletion of another two candidates (Tfap2c, Tead4) delayed cell polarization. Sequences of primers for generating dsRNA are listed in Table S4.

**Figure S4.**
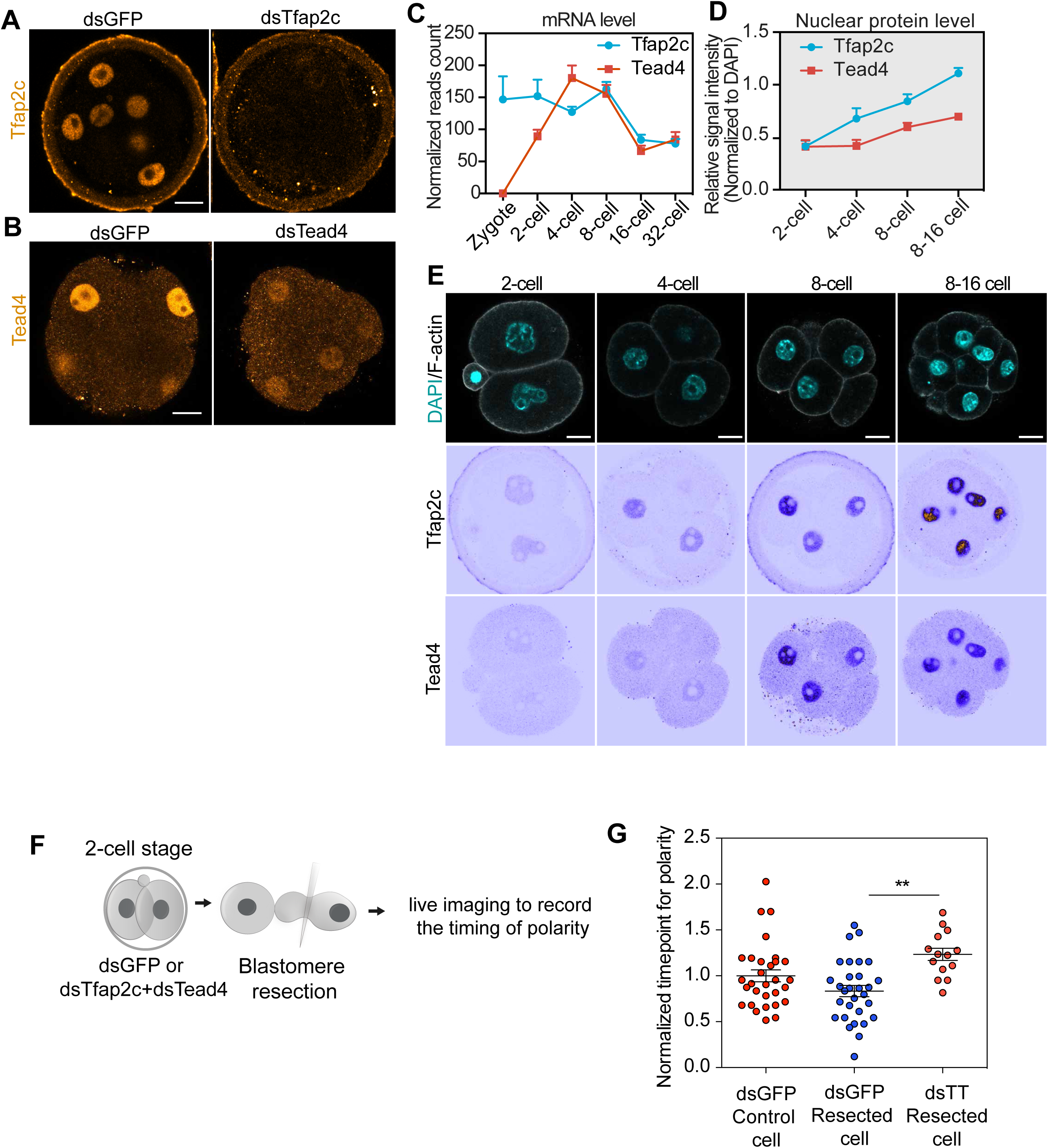
Tfap2c and Tead4 expression is required for cell polarization. (**A**) Representative images of embryos injected with dsGFP or dsTfap2c RNAs and immunostained to reveal Tfap2c at the 8-16 cell stage (N=30 embryos). (**B**) Representative images of embryos injected with dsGFP and dsTead4 RNAs and immunostained for Tead4 at the late 8-cell stage (N=15 embryos). N=2 independent experiments. (**C**) Expression profile of Tfap2c and Tead4 transcripts from zygote to the 32-cell stage. Data presented as means ± S.E.M. (**D**) Quantification of Tfap2c and Tead4 nuclear protein levels from the 2- to the 8-16 cell stage. Data presented as means ± S.E.M. N=7 embryos for 2-cell stage; N=15 embryos for 4-cell stage; N=26 embryos for early 8-cell stage; N=17 embryos for 8-16 cell stage. Note that although Tfap2c mRNA is maternally stored, the protein is only expressed after zygotic genome activation. N=3 independent experiments. (**E**) Expression profile of Tfap2c and Tead4 proteins from the 2-cell stage to the morula stage. (**F**) 2-cell stage embryos were injected with dsGFP or dsTfap2c+dsTead4, after which the cells were separated, and one blastomere was subjected to resection. The timing of cell polarization in each group was determined by live imaging. (**G**) Quantification of the timing of cell polarization in control or resected cells from embryos injected with dsGFP or dsTfap2c+dsTead4. Each dot represents one embryo. 10 embryos injected with dsTfap2c+dsTead4 failed to polarize throughout the live imaging and were therefore removed from quantification. **p<0.01, student’s t-test. Scale bars, 15µm.

**Figure S5.**
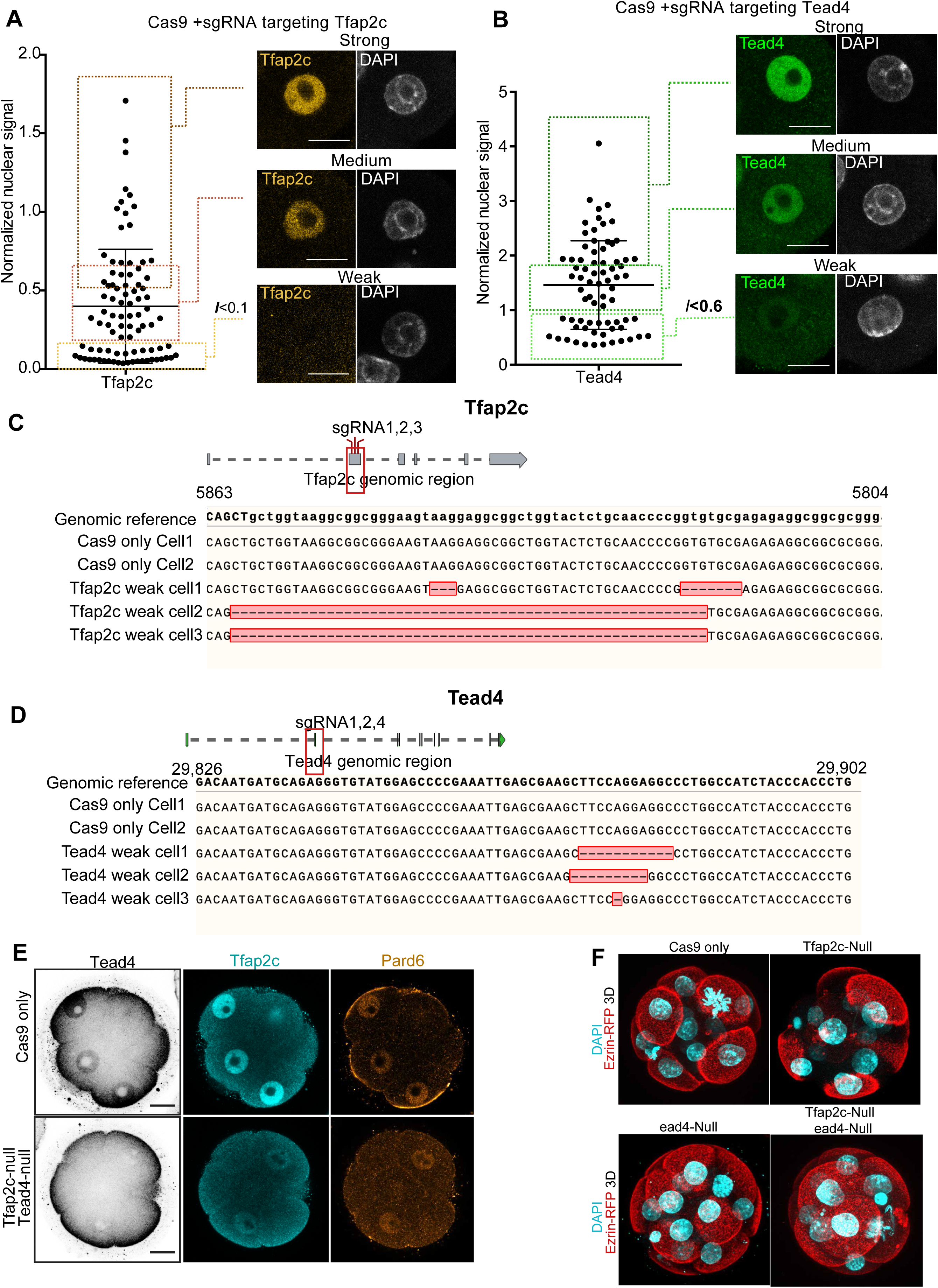
Genetic depletion of Tfap2c and Tead4 leads to the failure of cell polarization. (**A**) Signal intensity quantification in cells with strong, medium or low levels of Tfap2c (classification shown in squares). Representative images of cells expressing different levels of Tfap2c are shown on the right. Each dot represents a single cell (N=23 embryos). Normalization was performed by calculating the ratio of Tfap2c signal intensity over that of DAPI. A normalized ratio of less than 0.1 was considered to represent a “Tfap2c-null” cell. All cells are from embryos injected with Cas9 and gRNAs targeting Tfap2c were stained with anti-Tfap2c and DAPI at 8-16 cell stage. N=4 independent experiments. (**B**) Quantifications of the signal intensity for cells with strong, medium or low level of Tead4 (classification was shown in squares). Each dot represents a single cell (N=16 embryos). Representative images of cells expressing different levels of Tead4 are shown on the right. Each dot represents a single cell. Normalization was performed by calculating the ratio of Tead4 signal intensity over that of DAPI. A normalized ratio of less than 0.6 was considered to represent a “Tead4-null” cell. N=3 independent experiments. All cells are from embryos injected with Cas9 and sgRNAs targeting Tead4 and stained for Tead4 and DAPI at 8-16 cell stage. (**C**) DNA-sequencing was performed to determine the genotype of cells injected with Cas9 only (as a control) or cells injected with gRNA targeting Tfap2c. The cells with a low level of Tfap2c (criterion described in A) were sequenced. (**D**) DNA-sequencing to determine the genotype of cells injected with Cas9 only (as a control) or cells injected with gRNA targeting Tead4. The cells with a low level of Tead4 (criterion described in B) were sequenced. (**e**) Embryos expressing wild-type (injected with only Cas9 mRNA as control); single depletion of Tfap2c (injected with Cas9 and sgRNAs targeting only Tfap2c); single depletion of Tead4 (injected with Cas9 and sgRNAs targeting only Tead4 sgRNAs); double depletion of Tfap2c and Tead4 (injected with Cas9 and sgRNAs targeting both Tfap2c and Tead4) were analyzed at the 8-16 cell stage to reveal Tfap2c, Tead4 and Pard6. N=2 independent experiments. (**F**) Ezrin-RFP and DAPI localization in embryos expressing wild-type (injected with only Cas9 mRNA as control); single depletion of Tfap2c (injected with Cas9 and sgRNAs targeting only Tfap2c); single depletion of Tead4 (injected with Cas9 and sgRNAs targeting only Tead4 sgRNAs); double depletion of Tfap2c and Tead4 (injected with Cas9 and sgRNAs targeting both Tfap2c and Tead4). Maximum projection is shown in all images. Scale bars, 15µm.

**Figure S6.**
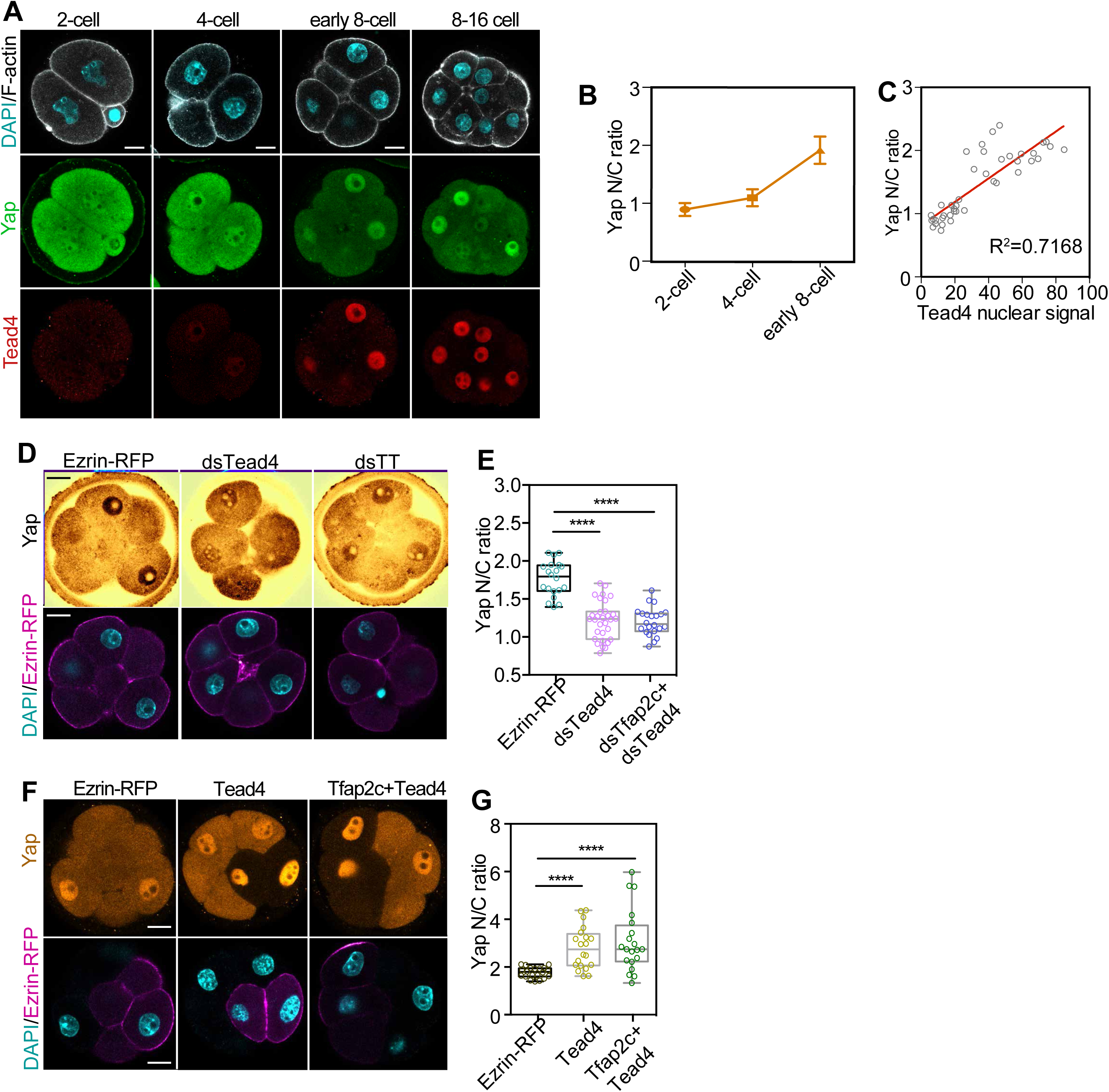
Tead4 expression drives Yap nuclear localization independently of the apical domain. (**A**) Expression profiles of Yap and Tead4 from the 2-cell to the 8-16 cell stage. Quantification is shown in **B**. (**B**) Quantification of Yap nuclear/cytoplasm (N/C) ratio from the 2- to the early 8-cell stage. Data presented as means ± S.E.M. N=10 cells from N=5 embryos for 2-cell stage, N=13 cells from N=4 embryos for 4-cell stage, N=21 cells from N=5 embryos for early 8-cell stage. N=2 independent experiments. (**C**) Positive correlation between Tead4 nuclear expression and Yap nuclear/cytoplasm ratio, each dot represents one analyzed cell. R^2^ = 0.7168, p<0.0001. (**D**) Embryos injected with Ezrin-RFP mRNA only (control), dsTead4 and dsTead4+Tfap2c RNAs and analyzed at the early 8-cell stage to reveal Yap localization. (**E**) Quantification of Yap N/C ratio in cells injected with Ezrin-RFP mRNA only, or co-injected with dsTead4, dsTfap2c+dsTead4 RNAs. Data shown as individual data points with Box and Whiskers plot (bottom: 25%; upper: 75%; line: median; whiskers: min to max). N=20 cells from N=4 embryos for Ezrin-RFP only; N=18 cells from N=3 embryos for dsTead4; N=27 cells from N=5 embryos for dsTfap2c+dsTead4. N=2 independent experiments. ****p<0.0001. One-way ANOVA test. (**F**) Embryos microinjected with Ezrin-RFP mRNA only, Tead4 and Tead4+Tfap2c mRNAs and analyzed for Yap localization at the early 8-cell stage. Arrows indicate cells overexpressing Tead4. (**G**) Quantification of Yap N/C ratio in cells microinjected with Ezrin-RFP mRNA only; with Tead4; or with Tfap2c+Tead4 mRNAs. Data shown as individual data points with Box and Whisker plots (lower: 25%; upper: 75%; line: median; whiskers: min to max). N=20 cells for each group, from N=3 embryos for Ezrin-RFP group; N=4 embryos for Tead4 group and N=5 embryos for Tfap2c+Tead4 group. N=2 independent experiments. ****p<0.0001, one-way ANOVA test. Tead4 knockdown decreases Yap nuclear localization and conversely Tead4 overexpression enhances Yap nuclear localization by the early 8-cell stage. Scale bars, 15µm.

**Figure S7.**
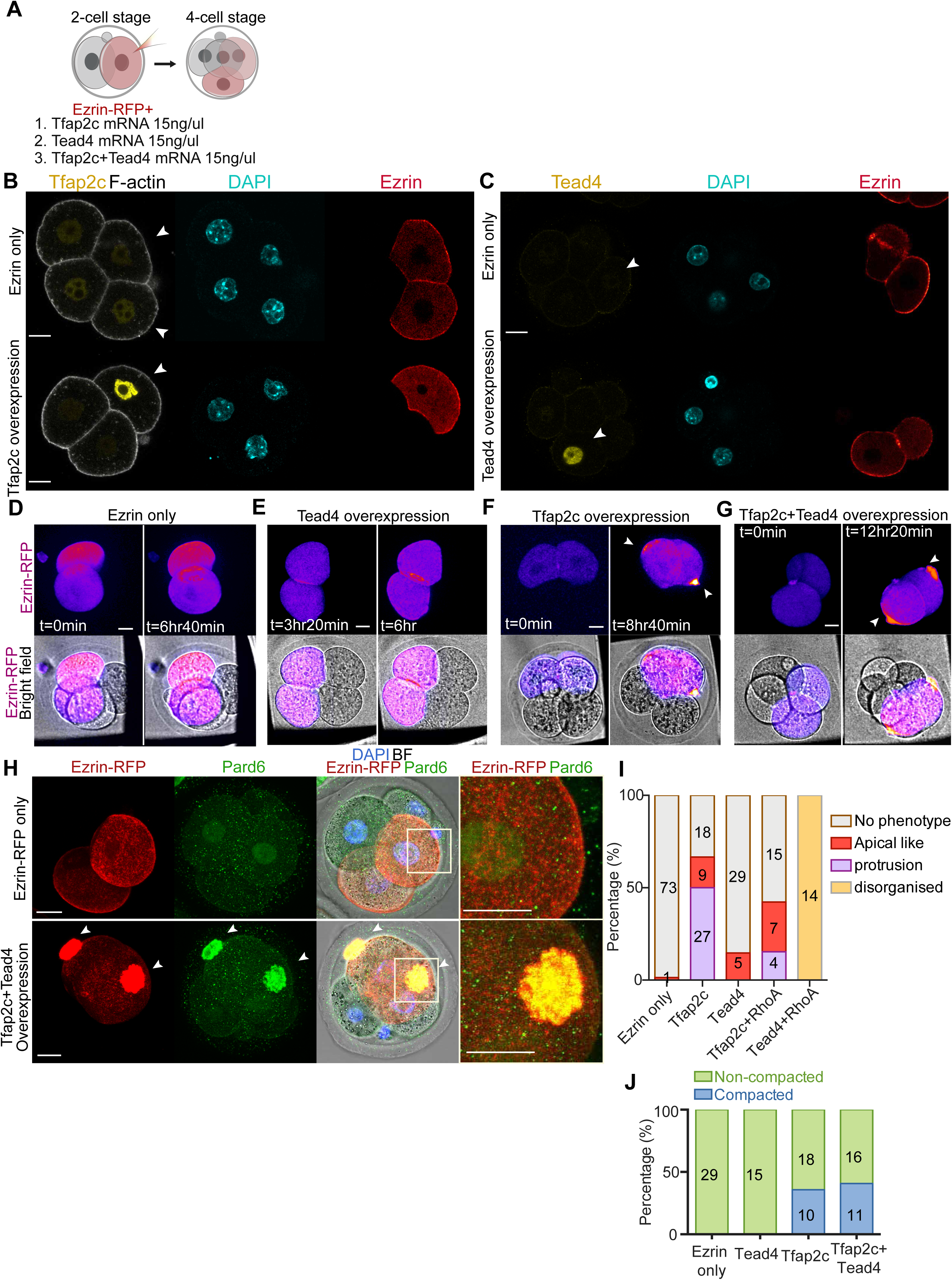
Advancing the activity of Tfap2c, Tead4 and RhoA-Q63L induces premature apical domain formation. Overexpression of Tfap2c and Tead4 induced cell protrusions enriched by apical proteins. (**A**) Schematic of experiment in which one 2-cell stage blastomere was injected with mRNA encoding Ezrin (as a control), Tfap2c, Tead4 or Tfap2c+Tead4. (**B**) Embryos injected with mRNA encoding Tfap2c and Ezrin-RFP (as an injection marker) in one blastomere at the 2-cell stage were analyzed at the 4-8 cell stage to reveal DAPI, F-actin, Tfap2c and Ezrin-RFP. N=7 embryos per each condition were examined. N=1 independent experiment. (**C**) Embryos injected with mRNA encoding Tead4 and Ezrin-RFP (as an injection marker) in one blastomere at the 2-cell stage were analyzed at the late 4-cell stage to reveal DAPI, F-actin, Tead4 and Ezrin-RFP. N=8 embryos for Ezrin only and N=14 embryos for Tead4 overexpression were examined. For both **B** and **C**, arrows indicate injected cells. N=2 independent experiments. (**D**-**G**) Representative images of embryos injected with Ezrin-RFP alone (**D**) or with Tead4 (**E**), Tfap2c (**F**) or Tead4+Tfap2c (**G**) mRNA at the late 4-cell stage. Arrows indicate cell protrusions induced by Tfap2c or Tfap2c+Tead4 overexpression. Quantifications are shown in **I** and Fig. 3C. (**H**) Embryos injected with Ezrin-RFP only (control), or co-injected with Tfap2c+Tead4 mRNAs in one of the two blastomeres at the 2-cell stage and analyzed at the 4-8 cell stage to reveal Pard6, Ezrin-RFP and DNA. Arrows indicate protrusions/apical domains. (**I**) Quantification of structures induced by different conditions. Data presented as a stacked bar graph where numbers in each bar indicate number of cells analyzed. N=29 embryos for Ezrin-RFP + LifeAct-GFP group, N=21 embryos for Tfap2c overexpression group; N=15 embryos for Tead4 overexpression group; N=13 embryos for Tfap2c+RhoA-Q63L overexpression group; N=43 embryos for Tfap2c+Tead4 overexpression group; N=4 independent experiments. (**J**) Quantification of compaction of cells expressing Ezrin (control), Tfap2c, Tead4, Tfap2c and Tead4 at the late 4-cell stage. Compaction was assessed based on the intercellular angle as previously described (Zhu et al., 2017). Numbers in each bar indicate the number of embryos analyzed. N=4 independent experiments. Scale bars, 15µm.

**Figure S8.**
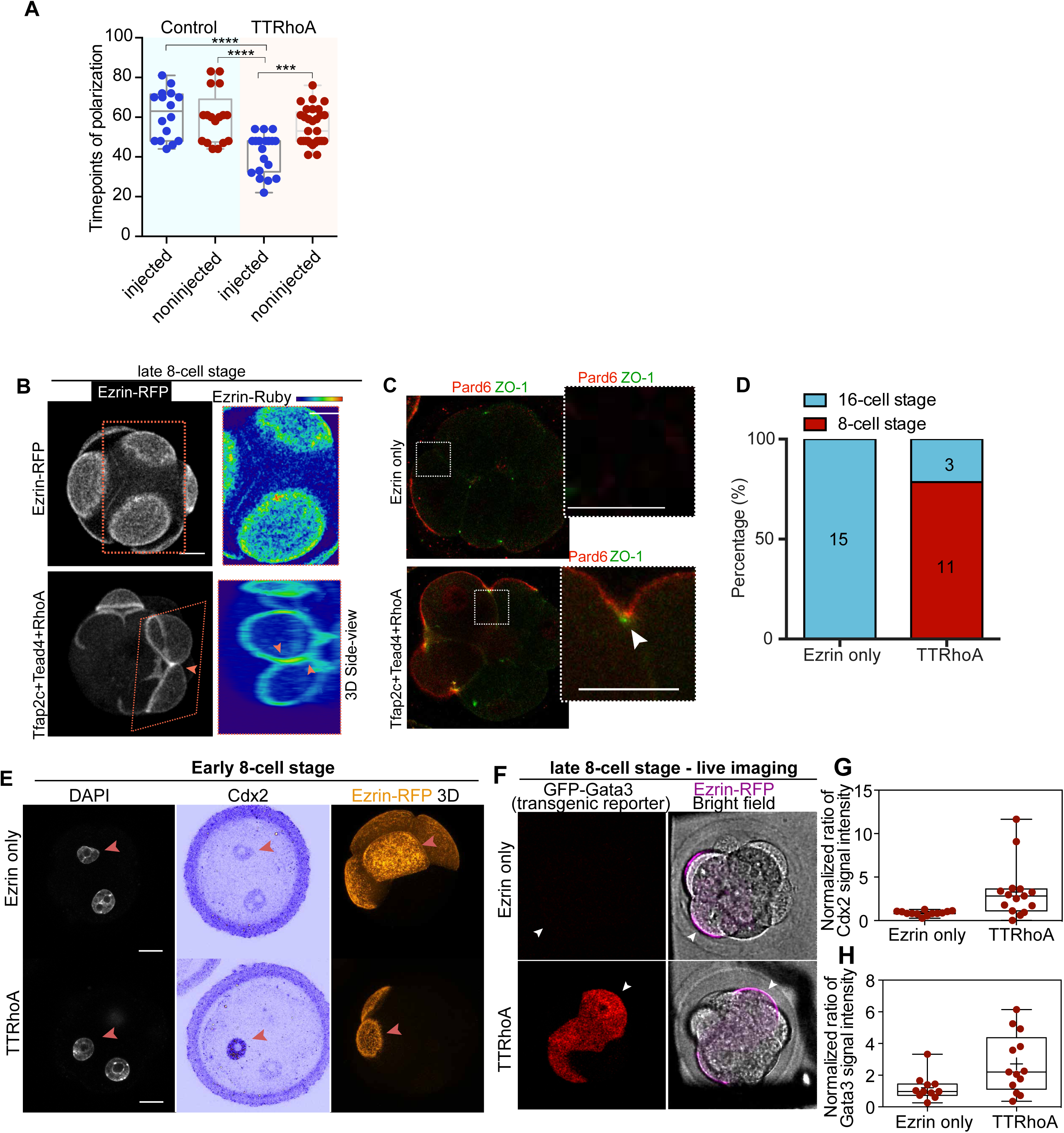
Premature expression of Tfap2c, Tead4, and activated RhoA is sufficient to advance the timing of polarization and differentiation. **(A)** Overview of the comparison of the timing of cell polarization in cells with or without overexpression of Tfap2c, Tead4 and RhoA in the same embryo or in cells of embryos with or without injection of a control marker (LifeAct-GFP). Timing of cell polarization was quantified from the timepoint at which the first cell of an embryo becomes polarized. ***p<0.001; ****p<0.0001, ordinary One-way ANOVA test. (**B**) Apical domain zippering in embryos expressing Tfap2c+Tead4+RhoA-Q63L at the late 8-cell stage (N=24 embryos). Pink and Yellow rectangular outlines indicate different magnified regions (below). Colored arrows indicate boundaries of apical domains in different cells. N=4 independent experiments. (**C**) Embryos expressing Ezrin-RFP (control) and Tfap2c+Tead4+RhoA-Q63L at the late 8-cell stage stained to reveal Pard6 and the tight junction marker, ZO-1 (N=8 embryos for Ezrin alone, N=12 embryos for Tfap2c+Tead4+RhoA-Q63L). Magnified regions are outlined. Arrows indicate the points where the apical domain and tight junction converge. N=2 independent experiments. (**D**) Quantification of the timing of zippering in adjacent blastomeres in control embryos or embryos overexpressing Tfap2c, Tead4, and RhoA-Q63L showing premature cell polarity at the 4-cell stage in the latter group. Data presented as a stacked bar graph where numbers in each bar indicate the number of cells analyzed. (**E**) Embryos expressing Ezrin-RFP (control) and Tfap2c+Tead4+RhoA-Q63L at the mid 8-cell stage stained to reveal DNA, Cdx2, and Pard6. Injected cells (arrows) express Ezrin-GFP (N=8 embryos). N=4 independent experiments. (**F**) Quantification of normalized Cdx2 expression in either cells injected with Ezrin-RFP or with Tfap2c+Tead4+RhoA-Q63L mRNAs as in **E**. The intensity of Cdx2 expression in cells of each group is normalized to the intensity of DAPI (DNA) in the same cell; injected cells were normalized against non-injected cells in the same embryo for each group. Each dot represents a single embryo. ****p<0.0001, Mann-Whitney test. Overexpression of Tfap2c+Tead4+RhoA-Q63L advanced the timing of apical domain formation and upregulated Cdx2 expression. Arrows indicate an injected cell. (**G**) GFP-Gata3 transgenic embryos were injected with Ezrin-RFP mRNA alone (as a control) or together with Tfap2c/Tead4/RhoA-Q63L and imaged at the late 8-cell stage for RFP and GFP to reveal Gata3 expression level. (**H**) Quantification of normalized GFP-Gata3 expression in cells injected with Ezrin-RFP or with Tfap2c+Tead4+RhoA-Q63L mRNAs as in **G**. The intensity of GFP expression in injected cells of each group is normalized to the intensity of non-injected cells. Each dot represents a single embryo. *p<0.05, Mann-Whitney test. Overexpression of Tfap2c+Tead4+RhoA-Q63L induced earlier Gata3 expression. N=2 independent experiments. Arrows indicate an injected cell. Scale bars, 15µm.

**Figure S9.**
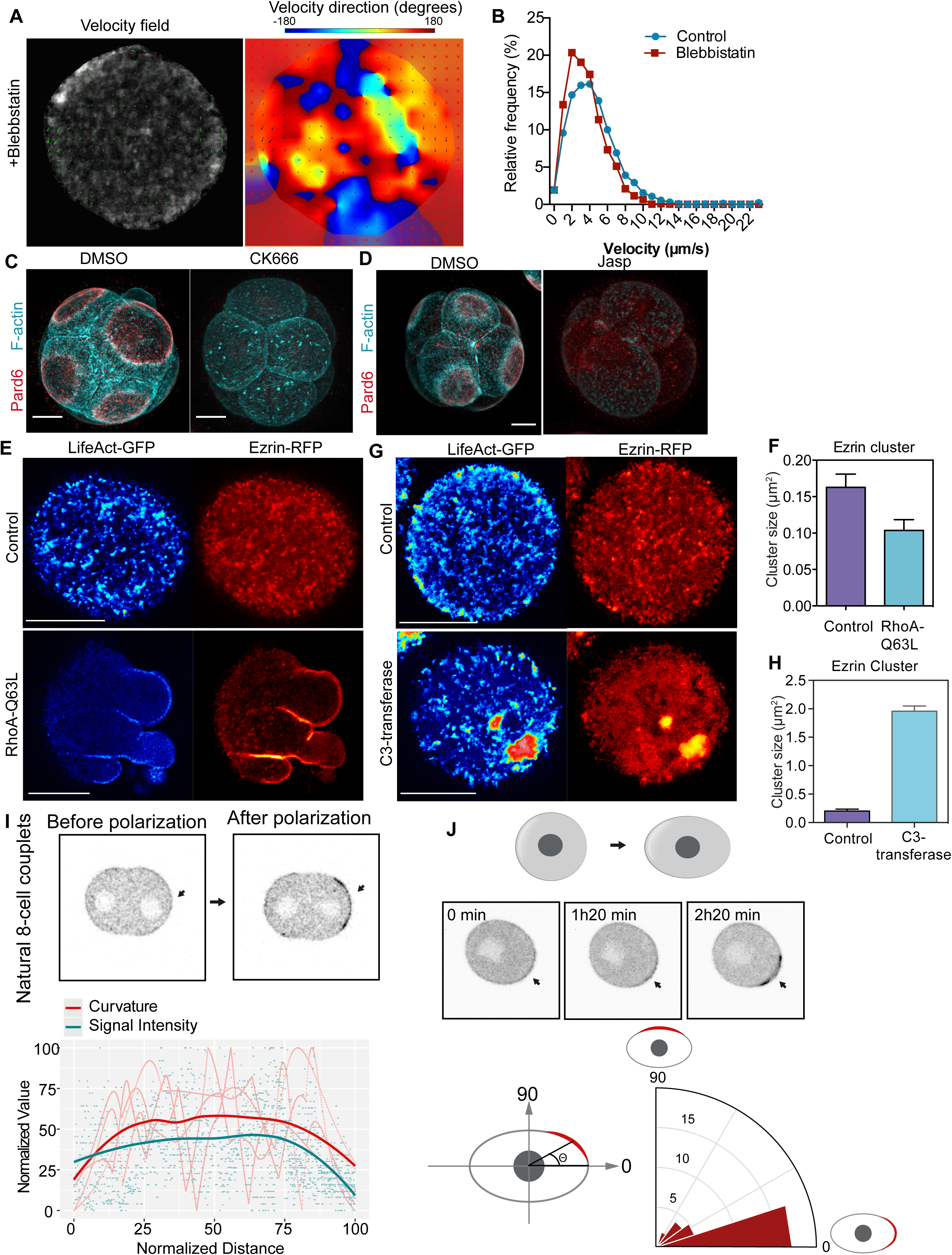
Actin turnover and RhoA signaling regulates the apical protein clustering and apical domain formation. (**A**) PIV analysis and vector velocity quantification of the cells treated with Blebbstatin. (**B**) Histogram showing vector velocity distribution of LifeAct-GFP flow in cells with or without Blebbstatin. (**C**) Representative images of embryos treated with DMSO or CK666 showing of Pard6 or F-actin. N=8 embryos examined per condition, N=2 experiments. (**D**) Representative images of embryos treated with DMSO or Jasp showing the localization of Pard6 or F-actin. N=10 embryos for control, and N=24 embryos for Jasp treated embryos examined, N=2 experiments. (**E**) Localization of LifeAct-GFP and Ezrin-RFP in the cells with or without overexpression of RhoA-Q63L. (**F**) Quantification of the size of Ezrin clusters in cells with or without overexpression of RhoA-Q63L. N=5 cells for each conditions, N=2 experiments. (**G**) Localization of LifeAct-GFP or Ezrin-RFP in embryos treated with water (as a control) or C3-transferase. (**H**) Quantification of the size of Ezrin clusters in cells treated with water or C3-transferase. N=5 cells from water group, N=4 cells from C3-transferase group. N=2 experiments. (**I**) Representative images and quantification of cell curvature along the cell-contact free surface of compacted 8-cell stage embryos. Arrows indicate the point of high curvature. To quantify the correlation and any influence of polarization on cell curvature, a time-lapse movie was run form the early 8-cell stage to the late 8-cell stage. The cell curvature was measured at the timepoint when the cells undergo compaction but are not polarized. The signal intensity of the apical domain was recorded when cells polarized. N= 4 cells from N=4 8-cell stage pairs were measured. N=2 experiments. (**J**) Representative images and quantifications of the correlation between the position of the apical domain and the long axis of the cell in elongated 8-cell stage cells. The early 8-cell stage cells were elongated (see methods) and a time-lapse movie was run to record the position of the apical domain after the elongation procedure. N=34 cells from N=2 experiments. Scale bars, 15µm.

**Figure S10.**
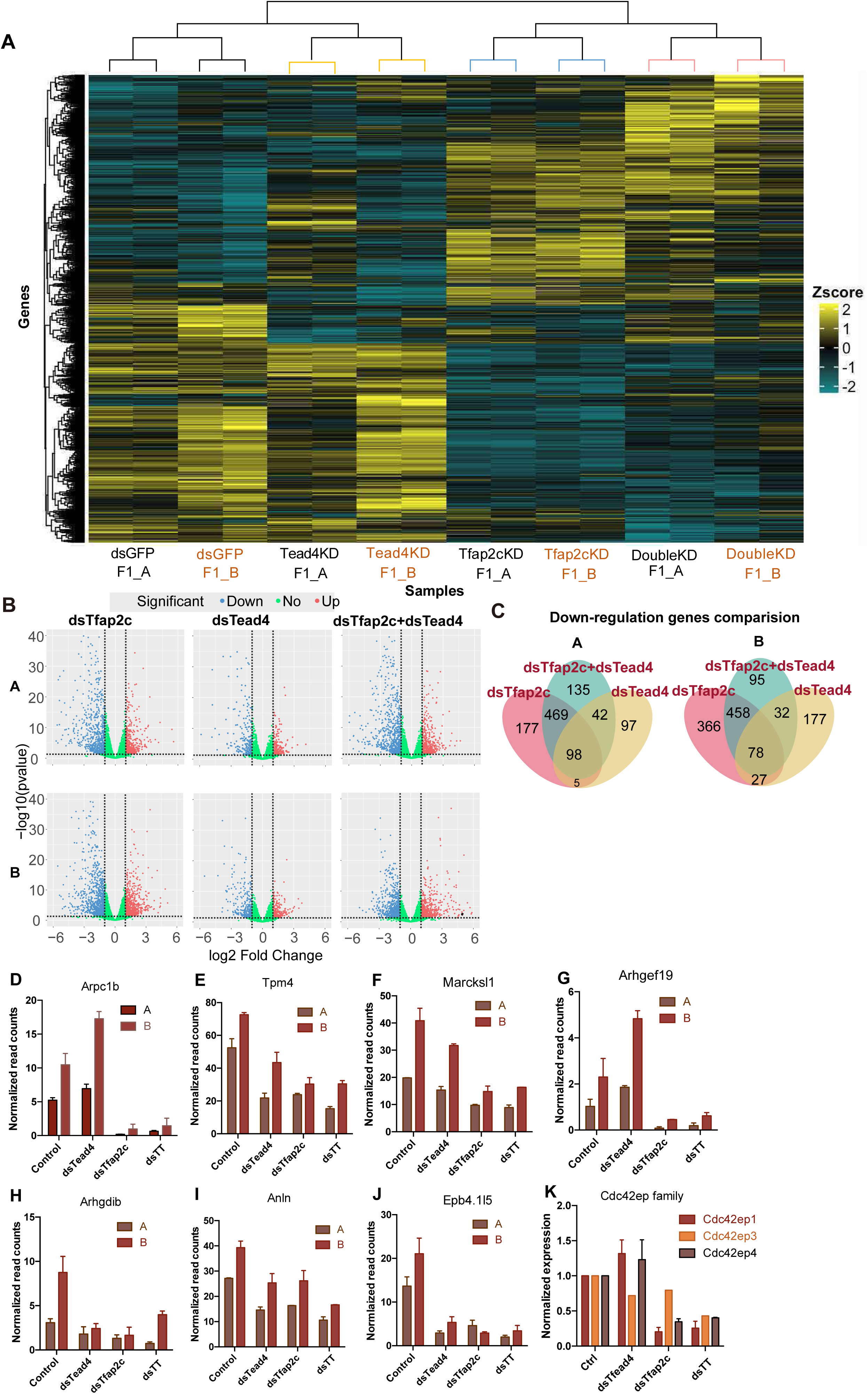
Tfap2c and Tead4 control gene expression at the early 8-cell stage. (**A**) Heatmap plot shows the expression profile of differentially expressed genes between embryos injected with dsGFP (as a control) and dsTfap2c, dsTead4 or dsTfap2c+dsTead4 in two different genetic backgrounds at the 8-cell stage. (**B**) Volcano Plots show the genes that were up- or down-regulated upon the depletion of Tfap2c, Tead4 or Tfap2c+Tead4 in different genetic backgrounds. For each group 2 or 3 replicates were analyzed and genes showing consistent up- or downregulated are shown. Dotted lines indicate the cut-off boundaries for significantly up- or downregulated genes. (**C**) Venn plot shows the number of genes downregulated in Tfap2c, Tead4 or Tfap2c+Tead4 knockdown groups in two distinct genetic backgrounds. (**D**-**J**) Expression levels of cell polarity regulators upon the depletion of Tead4, Tfap2c or Tead4+Tfap2c. A indicates the embryos obtained from the mating ♀C57BL/6J×CBA/J ♂C57BL/6J×DBA/2J; B indicates the embryos obtained from the mating♀C57BL/6J×DBA/2J **♂**C57BL/6J×DBA/2J. Data shown as mean ± S.E.M. (**K**) Normalized expression levels of various Cdc42ep family members upon the depletion of Tead4, Tfap2c or Tead4+Tfap2c. For all graphs, data are shown as mean ± S.E.M.

