## Supplementary information for "Transcriptional control of apical protein clustering drives *de novo* cell polarity establishment in the early mouse embryo"

**Table S1.** List of candidates that are selected from single cell RNA sequencing for the polarity regulators screen.

**Table S2.** List of transcription factors that are selected from ATAC-seq analysis for the polarity regulators screen.

**Table S3.** List of genes associated with actin regulation that show downregulation upon the depletion of Tfap2c and/or Tead4.

**Table S4.** Primer sequences for sgRNA, dsRNA preparation and for real-time qPCR experiments.

**Movie S1.** Time-lapse imaging of control sister cell and resected sister cell expressing Ezrin-RFP developing from 4-cell stage to blastocyst stage from experiment illustrated in Figure 1e. Scale bars, 15µm. Time interval, 1 h. Arrows indicate the apical domains.

**Movie S2.** Time-lapse imaging of control sister pair or resected sister pair expressing Ezrin-RFP developing from 4-cell stage to blastocyst stage from experiment illustrated in Supplementary Fig. 4a. Scale bars, 15µm. Time interval, 1 h1. Arrows indicate the apical domains.

**Movie S3.** Time-lapse imaging of the natural apical domains formation process in a control embryo expressing Ezrin-RFP during the 4- to 8-cell stage development. Scale bar, 15µm. Time interval, 22min.

**Movie S4.** Time-lapse imaging of the membrane protrusions formation induced by Tead4 and Tfpa2c overexpression at the late 4-cell stage. Scale bar, 15µm. Time interval, 22min. Arrows indicate the membrane protrusions.

**Movie S5.** Time-lapse imaging of the membrane disorganization induced by RhoA-Q63L overexpression at the late 4-cell stage. Scale bar, 15µm. Time interval, 22min.

**Movie S6.** Time-lapse imaging of the apical domains formation induced by Tead4, Tfap2c and RhoA-Q63L overexpression. Scale bar, 15µm. Time interval, 22min. Arrows indicate the apical domain.

**Movie S7.** Natural embryo injected with Ezrin-RFP mRNA in whole embryo, and LifeAct-GFP mRNA in half of the embryo at the 2-cell stage, developing from 4- to late morula stage. Both LifeAct-GFP injected or non-injected cells develop the apical domain at the late 8-cell stage. Scale bar, 15µm. Time interval, 30min.

**Movie S8.** Embryo injected with Ezrin-RFP mRNA in whole embryo, and Tead4, Tfap2c, RhoA-Q63L and LifeAct-GFP mRNA in half of the embryo, developing from 4- to late morula stage. Tead4, Tfap2c, RhoA-Q63L and LifeAct-GFP mRNA injected cells establish the apical domains at the late 4-cell stage, whereas non-injected cells develop the apical domain at the late 8-cell stage. Scale bar, 15µm. Time interval, 30 min.

**Movie S9.** Gata3-GFP transgenic embryo injected with Ezrin-RFP mRNA in half of the embryos developing from 8- to the blastocyst stage. Gata3-GFP signal appears from the 16-cell stage in the control embryo. Scale bar, 15µm. Time interval, 30min.

**Movie S10.** Gata3-GFP transgenic embryo injected with Ezrin-RFP and Tead4, Tfap2c and RhoA-Q63L mRNA in half of the embryos developing from 8- to the blastocyst stage. Gata3-GFP signal is upregulated at the late 8-cell stage in the overexpressed cells of the embryo. Scale bar, 15µm. Time interval, 30min.

**Movie S11.** Natural embryo injected with Ezrin-RFP (whole embryo) and LifeAct-GFP (half of the embryo) mRNA imaged from mid 8-cell stage to the late 8-cell stage. Scale bar, 15µm. Time interval, 6min.

**Movie S12.** Natural embryo injected with Ezrin-RFP and LifeAct-GFP mRNA imaged from mid 8-cell to late 8-cell stage with short time-interval to reveal Ezrin and Actin dynamics. Scale bar, 15µm. Time interval, 1min.

**Movie S13.** Natural embryo injected with Ezrin-RFP and LifeAct-GFP mRNA imaged at the mid 8-cell stage prior to the development of the apical domain. Scale bars, 15µm. Time interval, 4s.

**Movie S14.** PIV analysis of the natural embryo injected with LifeAct-GFP mRNA imaged at the mid 8-cell stage prior to the development of the apical domain.

**Movie S15.** Embryos injected with Ezrin-RFP and LifeAct-GFP mRNA were treated with Blebbstatin and imaged at the mid 8-cell stage prior to the development of the apical domain. Scale bars, 15µm. Time interval, 4s.
